# Human gloss perception reproduced by tiny neural networks

**DOI:** 10.1101/2025.05.09.653112

**Authors:** Takuma Morimoto, Arash Akbarinia, Katherine R. Storrs, Jacob R. Cheeseman, Hannah E. Smithson, Karl R. Gegenfurtner, Roland W. Fleming

## Abstract

A key goal of visual neuroscience is to explain how our brains infer object properties like colour, curvature, or gloss. Here, we used machine learning to identify computations underlying human gloss judgments from object images—traditionally considered a challenging inference. Thousands of object images of varied shapes were rendered with a Ward reflectance model under diverse lighting and viewpoints, and we crowdsourced gloss estimates for each image. Curiously, observers’ judgments were highly consistent with one another, yet systematically deviated from reality. We compared these responses with neural networks trained either to estimate physical reflectance (“ground-truth networks”) or to reproduce human judgments (“human-like networks”). By reducing network size, we identified the minimum computations needed for each objective. While estimating physical reflectance required deep networks, shallow networks—with as few as three layers—accurately replicated human judgments. Remarkably, even a miniscule ‘network’ with just a single filter could predict human judgments better than the best ground-truth network. The human-like networks also captured known gloss illusions outside the training range. These results suggest human gloss judgments rely on simpler, more general-purpose computations than previously thought, and demonstrate the power of using ‘tiny’ but interpretable neural networks to uncover functional computations in human brains.

## Introduction

In daily life, we encounter materials with a wide range of qualities, like metal, cloth, or plastic [1]. In a brief glance, we can judge their colour, lightness, shape, glossiness, translucency and other appearance characteristics. We can infer associated physical properties, such as weight, softness, or roughness, and even their states, such as whether they are wet or dry, clean or dirty, solid or melting [2]. Humans use visually sampled information about materials to guide our behaviors, e.g. lifting heavy objects slowly or plucking ripe berries gently.

Among the various material properties that we can perceive visually, gloss is the most extensively studied [3]. The term *gloss* refers to the subjective impression of shine or lustre, arising from mirror-like surface reflections (see **Supplementary Information, SI** for details). Gloss estimation is widely considered a quintessential, challenging perceptual inference [3–5] as it involves not only distinguishing highlights and reflections from other features (e.g., bumps or surface markings) but also pooling and interpreting the reflections to arrive at a global estimate of the surface’s reflectance. Past studies have proposed various cues the visual system may use to determine the glossiness of an object’s surface, including the skewness of the luminance histogram [6–9], the luminance gradient [10], the standard deviation of luminance over a surface [11], and various image metrics derived from specular reflection patterns [12,13]. One significant challenge is that an object with specific material properties can differ substantially in visual appearance, depending on factors such as shape, lighting, and viewpoint. To characterise how the visual system addresses this challenge, many studies have measured gloss perception across a wide variety of viewing conditions [14–30]. Yet, no single model fully captures human gloss perception across these diverse factors.

In the present study, we aim to identify biologically plausible, *image-computable* models that can account for human gloss judgments across a broad set of images. Our focus is specifically on understanding the perceptual computations that support judgments of gloss *from images*. While this is not equivalent to gloss perception in real-world, dynamic viewing, image-based judgments provide a tractable and meaningful way to investigate the visual information and representations that correlate with perceived gloss.

### ‘Tiny’ neural networks as interpretable data-driven models

One persistent challenge for traditional cue-based approaches is that researchers must somehow invent candidate visual cues in advance to evaluate their efficacy. This can be a hurdle when the cues the visual system relies on may be too complex or abstract to formulate. Recent studies have taken a data-driven approach, seeking deep learning models that mirror the pattern of human gloss judgments [31,32]. Our approach differs fundamentally by aiming to create human-like convolutional neural networks (CNNs) by directly training them on human perceptual data. We gathered gloss judgments from hundreds of human observers for thousands of object images in an online experiment. Through optimization, the CNNs spontaneously acquired a set of filters useful for reproducing human-like gloss judgments, without manual feature selection. This allows exploration of a much larger search space, not limited by the experimenter’s imagination. We then systematically adjusted the depth of the CNNs to understand the computational power required to predict human gloss judgments, aiming to make them as shallow as possible to increase interpretability [33–35]. We then interrogated the mechanisms that emerged within the networks. This allows for a data-driven, hypothesis-free approach to discovering the computations underlying human perception across diverse viewing conditions [36].

**Figure 1** summarizes our approach. We generated 3,888 images from the full combination of 36 lighting environments, 36 object geometries, and 3 viewpoints. Using an online crowd-sourced experiment, each image was assigned a perceived gloss value based on human gloss judgments (‘human labels’), first averaged across two repetitions per observer and then averaged across at least three observers. The observer’s task was to adjust the specular reflectance parameter of a reference object until the test and reference objects appeared to have the same glossiness. The quality of the data was confirmed through laboratory-based validation experiments (see **Supplementary Information**). We then trained two sets of CNNs with various depths: one on images with human labels (‘human-like networks’) and another on images with physical ground-truth labels (‘ground-truth networks’). To ensure that the networks learned only the gloss computation—rather than tasks such as image segmentation—we trained them on images where the object was shown against a uniform gray background (sRGB = [127, 127, 127]). Finally, we compared the performance of the two sets of CNNs and examined the internal computations of the human-like networks to gain insights into mechanisms that reproduce human gloss judgements.

**Figure 1.**
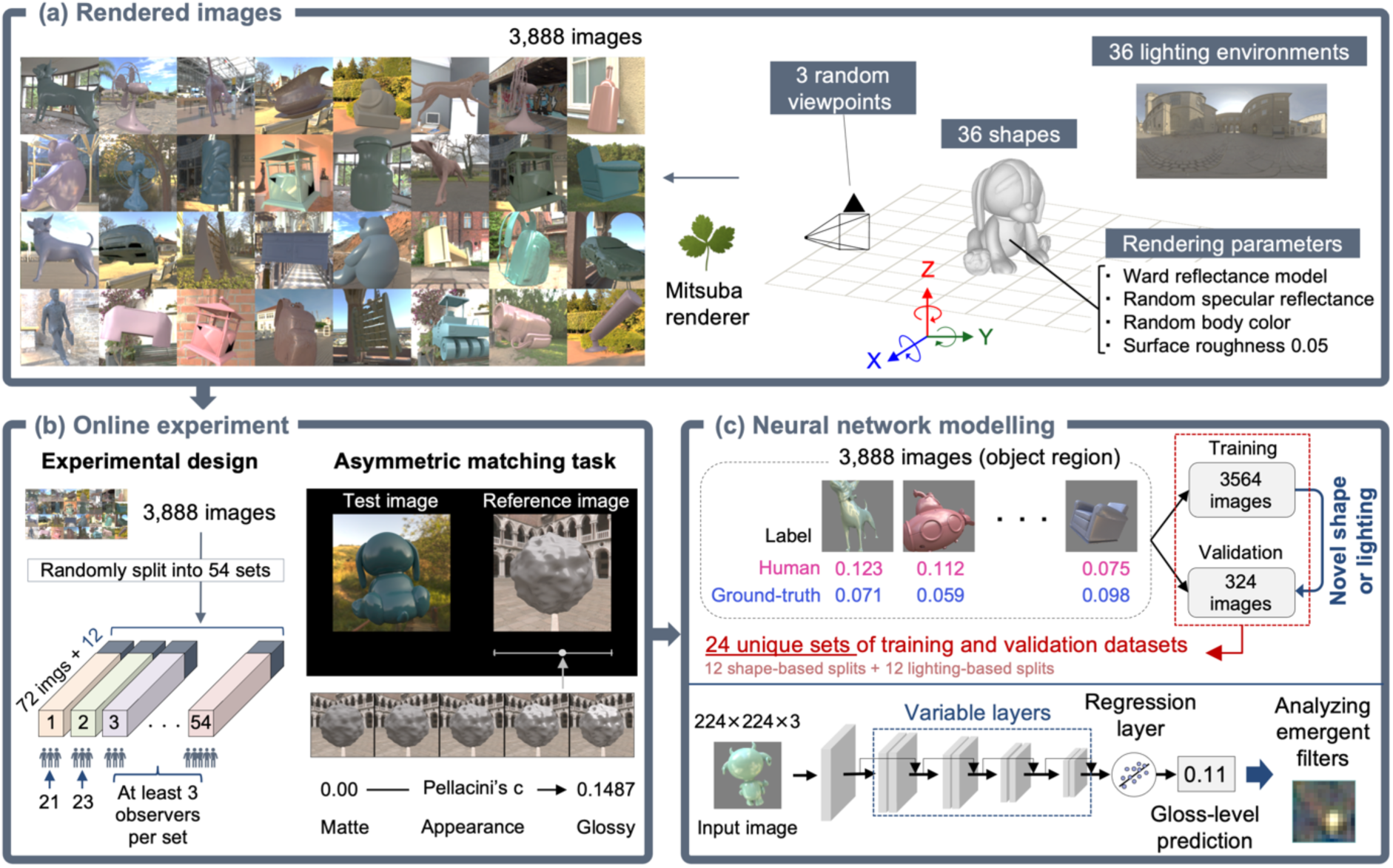
Schematic of our approach. **a** The scenes contained an object with a glossy surface (Ward reflectance model [37, 38], surface roughness fixed at 0.05). Random specular reflectance values and body colors were applied. The full set of 3,888 images was rendered by combining 36 shapes, 36 lighting environments, and 3 random viewpoints per shape-lighting pair (36 × 36 × 3). Images were generated in the CIE XYZ (1931) color space, converted to linear sRGB, and then displayed using standard sRGB gamma correction (γ = 2.4). **b Left**: Design of online experiment. The 3,888 images were randomly divided into 54 sets of 72 images each, plus 12 additional images that were shared across sets and rated by all online observers. These shared images, taken from our previous study (see Figure 5a in [30]), were used solely to assess inter-observer variability and were not included in the main set of 3,888 images or used for training the network models. Two image sets were also tested in a lab-based experiment (see **Supplementary Information**), to validate the online data quality. At least three observers were recruited for each set. **Right:** stimulus configuration for the asymmetric gloss matching task. By moving a slider, observers adjusted the gloss (i.e., Pellacini’s c [39]) of the reference object to match the perceived gloss level of the two objects. **c** For cross-validation, the 3,888 images were split such that each validation set consisted of a block of either three novel shapes (3 shapes × 36 lighting environments × 3 viewpoints = 324 images) or three novel lighting environments (36 shapes × 3 lighting environments × 3 viewpoints = 324 images), which were excluded from the corresponding training set. There were 12 such shape-based splits and 12 lighting-based splits, resulting in 24 unique, non-overlapping combinations of training (3,564 images) and validation (324 images) datasets. We trained CNNs with varying numbers of intermediate layers using images labeled by human gloss judgments (‘human-like networks’) or by physical ground-truth labels (‘ground-truth networks’), where each ‘label’ consisted of a continuous value of perceived gloss or physical specular reflectance as captured by Pellacini’s c. Gloss levels predicted by the networks were compared against human responses. The trained networks were analyzed to understand the computational mechanisms that emerged within them.

Our findings suggest that relatively shallow CNN architectures with as few as three convolutional layers are sufficient to predict human gloss judgments approximately as well as individual human judgments predict one another, while inferring the physical ground-truth requires far more complex computations for equivalent accuracy. Indeed, we find that even a single convolutional kernel can predict human perception better than the best of the ground-truth networks. Analysis of the emergent filters in the human-like networks showed that a filter resembling a combination of a Gaussian blob and diagonal ridge-like features can effectively extract image features predictive of human gloss judgments. This insight indicates that what seems to be a complex visual task, such as judging material properties, may be resolved through a set of simple computations that are also used for other visual tasks.

## Results

### Perceptual experiments

Each gray circle in **Figure 2a** represents one of the 3,888 test images. The axes display ground-truth specular reflectance values and observer settings (averaged over observers). Notably, the average observer settings significantly deviate from the ground-truth reflectance of the objects (Pearson’s *r*(3,888) = 0.52, *p* = 4.0×10^-272^). The histogram of Pearson’s correlation coefficients for each of the 295 observers shows a median correlation value of 0.46 (IQR = 0.40–0.54), with no observer exceeding 0.75 (**Figure 2b**). This indicates that gloss judgements deviated from ground truth in informative ways, likely due to the broader range of viewing conditions than in many earlier studies: by testing a wider range of conditions we were able to identify more cases where humans ‘misperceive’ the specular reflectance.

**Figure 2.**
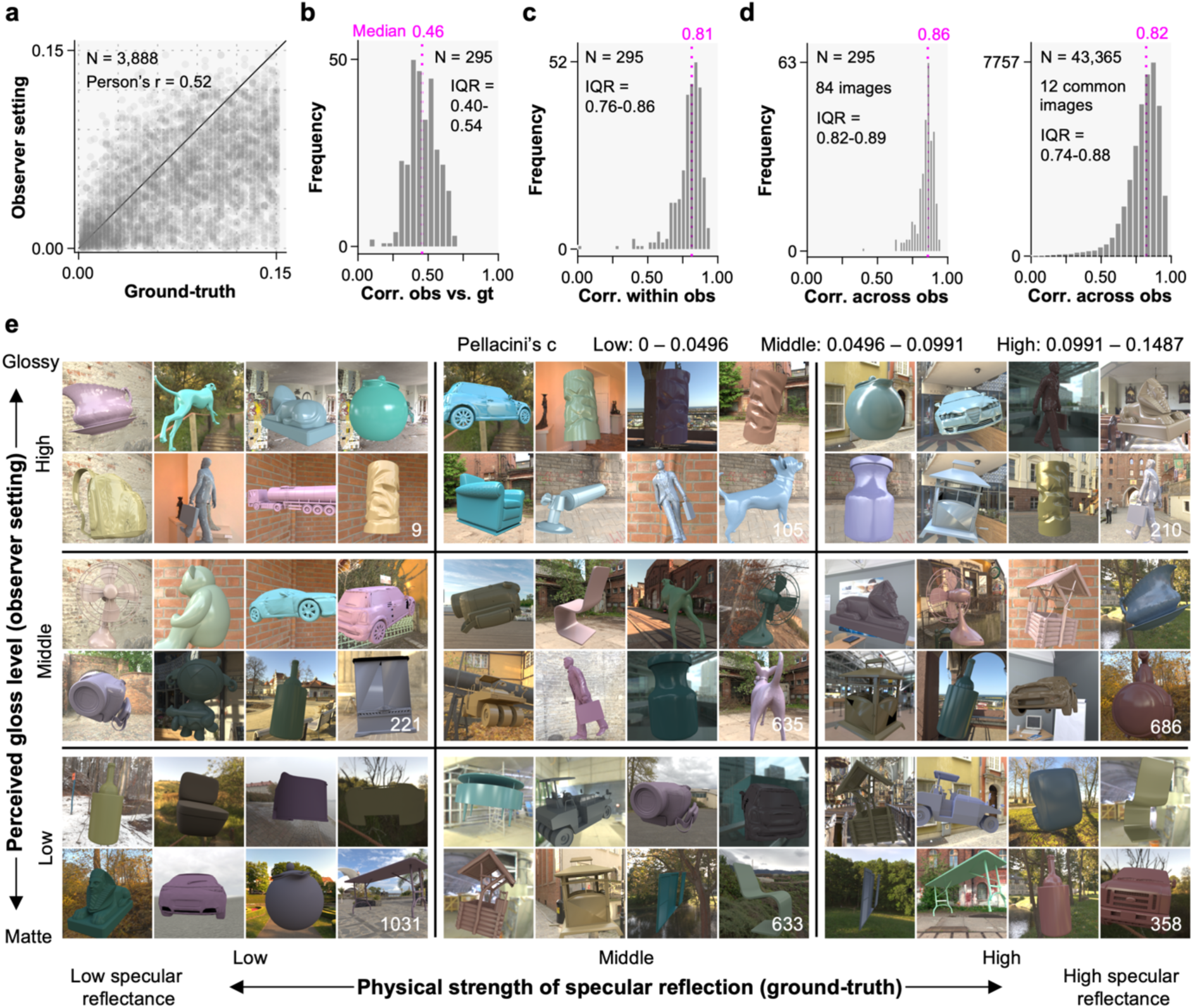
Summary of behavioural results. **a** Each data point shows a specific stimulus image, and the plot includes data on all 3,888 images. The horizontal and vertical axes show physical ground-truth reflectance values and observer settings (perceived gloss) averaged over observers. **b** Histogram of human vs. ground-truth correlation. The plot shows the distribution of Pearson’s correlation coefficients between each observer’s settings and the corresponding ground-truth values. **c** Histogram of within-observer correlations. For each observer, Pearson’s correlation coefficient was computed between session 1 and session 2 to quantify the intra-observer variability. **d** Histogram of correlation across observers. For the left plot, the correlation was computed between one observer and the average across all other observers across 72 images unshared across image sets. For the right plot, the correlation was computed over 12 images for every pair of observers (i.e. (295×294)/2 = 43,365 pairs). **e** Each grid shows 8 example images. The horizontal and vertical positions in the grid show the physical specular reflectance assigned to each object and observer settings (perceived gloss level). The number of images in each grid is shown at the right bottom of each grid.

However, crucially these gloss judgments were highly consistent both within and across observers (**Figures 2c** and **2d**). Intra-observer correlations were calculated between sessions 1 and 2 for each observer over 84 images (**Figure 2c**). Each inter-observer correlation was computed over 84 images between one observer and the average of the rest who judged the same set (left plot in **Figure 2d**), or between one observer and another for all possible observer pairs (295*294*0.5 = 43,365 pairs), over 12 common images (right plot). This confirms that the large deviation from the physical ground-truth was not due to random variation in observer judgments. On the contrary, observers made highly systematic and consistent patterns of agreement and disagreement with physical reflectance in gloss estimation (inter-observer correlation of 0.86 and 0.82; **Figure 2d**). A separate, lab-based experiment validated that the online data are of comparable quality (SI, Figure S6).

Each grid in **Figure 2e** shows eight example images in each pair of physical ground-truth and observer settings (low, mid and high). Images in the diagonal arrays (low-low, mid-mid, high-high) display objects where ground-truth and human judgments agree. Images in the top left and bottom right grids (low-high, high-low) highlight interesting cases where human judgments and physical ground-truth disagree substantially. Note that objects with low specular reflectance still have nonzero reflectance and can therefore produce visible highlights. Such images effectively provide different objectives for networks trained on physical ground-truth versus human judgments. Although human judgments do correlate somewhat with ground-truth, there is also sufficient consistent deviation from physical ground-truth to allow us to capture the key idiosyncratic computational signatures of human gloss perception.

### Computational models

**Figure 3** compares observers and candidate models based on their correlation to the physical ground-truth (x-axis) and human judgments (y-axis). The upper left half of the plot (pale blue region) indicates human-like models. The other half (pale pink region) indicates models that correlate more closely with the physical ground-truth. Each dark green circle represents an individual observer, where the y-axis shows the correlation to the average of the rest of the observers who judged the same 72 images. Thus, we are seeking models positioned close to this distribution of green circles in this plot.

**Figure 3.**
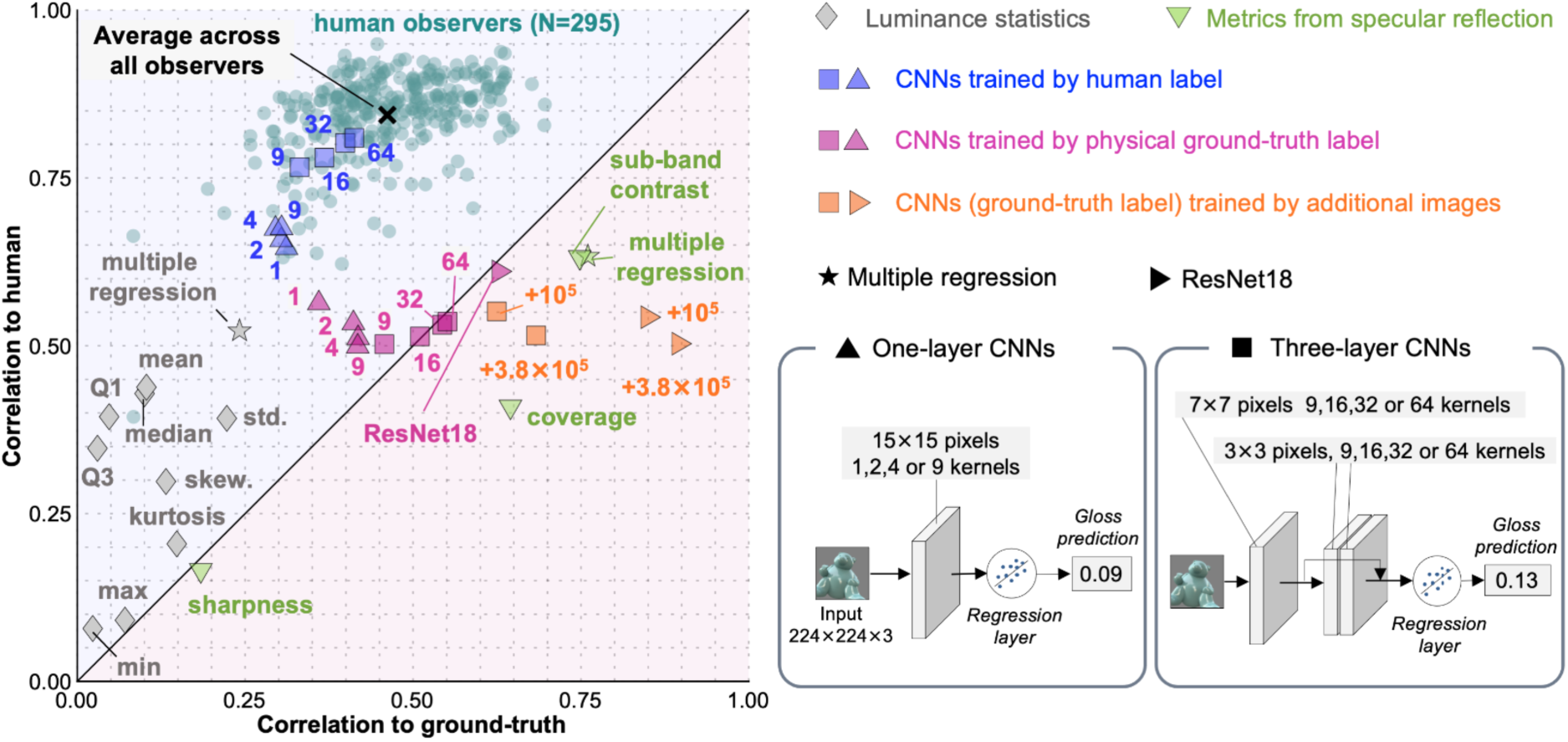
Summary of computational model behaviours. Dark green circles show individual human observers (N=295), where the x-axis and the y-axis show the correlation over 74 images to the physical ground-truth and the correlation to the average of the rest of the observers, respectively. Other data points show candidate computational models. Gray diamonds show low level luminance image statistics models, and the gray star shows a multiple regression model based on the luminance statistics (best fit to human data). Light green downward triangles show models based on statistics computed from specular reflection images (sharpness, coverage, and sub-band contrast), and the light green star symbol shows a multiple regression based on the specular metrics (best fit to human data). Triangles and squares show one-layer models with different kernel numbers (1, 2, 4 and 9; 9, 16, 32 and 64), where blue symbols refer to the networks trained on images labeled by observer judgments and pink symbols show networks trained on images labeled by physical ground-truth labels. The magenta rightward triangle shows ResNet18 trained on physical ground-truth labels. Orange symbols represent CNNs trained with additional images, generated using novel shapes and lighting conditions not included in the 3,888 images. Orange squares correspond to the three-layer model with 64 filters, and orange rightward triangles show ResNet18. The orange numbers on the plot show the number of additional images used.

Gray diamonds show models based on statistics of luminance distributions withing the image (mean, median, first quartile, third quartile, minimum, maximum, standard deviation, skewness, kurtosis), which have previously been implicated in gloss perception [40]. Yet here, we find they are positioned far from the human distribution. When these cues are combined using multiple linear regression fit to predict human responses, the correlation with human responses increases (indicated by the gray star symbol). Light green downward triangles show models that use sharpness, coverage and sub-band contrast computed from specular reflection images (a set of cues that summarize spatial variations in luminance, and that have been proposed to play a key role in gloss perception) [12]. These models correlate more with physical ground-truth than with human observers, indicating that they do not exhibit the characteristic errors seen in human judgments. Combining these cues yielded no improvement (the light-green star symbol) compared to the sub-band contrast model.

The blue triangles and squares show CNNs trained on human responses with one and three convolutional layers, respectively. The number next to each symbol shows the number of kernels in each convolutional layer. As expected, the deeper networks, such as three-layer with 64 kernels, came closer to approximating the mean glossiness judgments across all observers (black cross). Yet it is extremely intriguing and noteworthy that CNNs with only one kernel (single-kernel model) come close to the edge of the human distribution, with a mean correlation coefficient of 0.65 (SD = 0.064) across 24 independently trained models from 24-fold cross-validation. Given the correlation coefficient across observers was around 0.85, these very light models reach 75.3% of this performance ceiling and indeed approximate human perception better than any of the models trained on physical ground-truth labels that we tested.

To further contrast the human-trained models with an inverse-optics framework, we also trained networks on physical ground-truth labels and additional images, assuming that with sufficient training a supervised neural network can approximate a near-optimal observer at inferring specular reflectance from the images in our training set. Note that ‘near-optimal observer’ does not imply perfect performance, as some stimuli can be fundamentally ambiguous in that they contain insufficient information to infer reflectance accurately. Comparing human-like and physically trained networks in terms of their architectural complexity and training data requirements helps reveal how the underlying computations differ between recovering physical surface properties and predicting human perceptual judgments. Put another way, it allows us to test the extent to which human behaviour resembles a ‘near-optimal’ observer. Interestingly, we find that it does not.

In contrast to the CNNs trained on human labels, the networks trained on physical ground-truth labels, shown by magenta symbols, struggle to achieve high performance in inferring physical reflectance. Although trained to recover physical labels, some physical ground-truth networks are positioned near human-like networks, and even the three-layer models cluster around the diagonal unity line in the plot. A more complex architecture, such as ResNet18 (magenta rightward triangle), is also located near the unity line. Using additional training images rendered using novel lighting environments and object shapes, the network’s ability to predict ground-truth substantially improved, as shown by orange symbols. Three-layer models with 64 kernels (orange squares) still show a limited correlation below 0.70, even with an additional 3.8×10^5^ images. A ResNet18 trained with these additional images achieves a correlation of around 0.9 with the physical ground-truth, highlighting the computational challenges involved in estimating physical ground-truth. Importantly, however, its correlation with human responses remains below 0.5. The intriguing aspect of this observation is that it suggests that human strategies for estimating material properties such as specular reflectance are unlikely to rely on complex computations that aim to recover physical parameters, as suggested by inverse optics approaches. Instead, it appears that humans use relatively simple computations to intuit glossiness. Having established that lean neural network models can be trained to approximate human responses, we next sought to analyse the inner workings of these networks as a means to gain insights into the computations underlying human gloss judgements.

### Analysis of single-kernel models

As shown in **Figure 4a**, the single-kernel model first applies a 15×15×3 convolution to the input image, followed by max pooling and the addition of a bias term to predict a gloss level. To evaluate the variability of the emergent kernel, we analyzed 24 networks trained as part of the cross-validation process.

**Figure 4.**
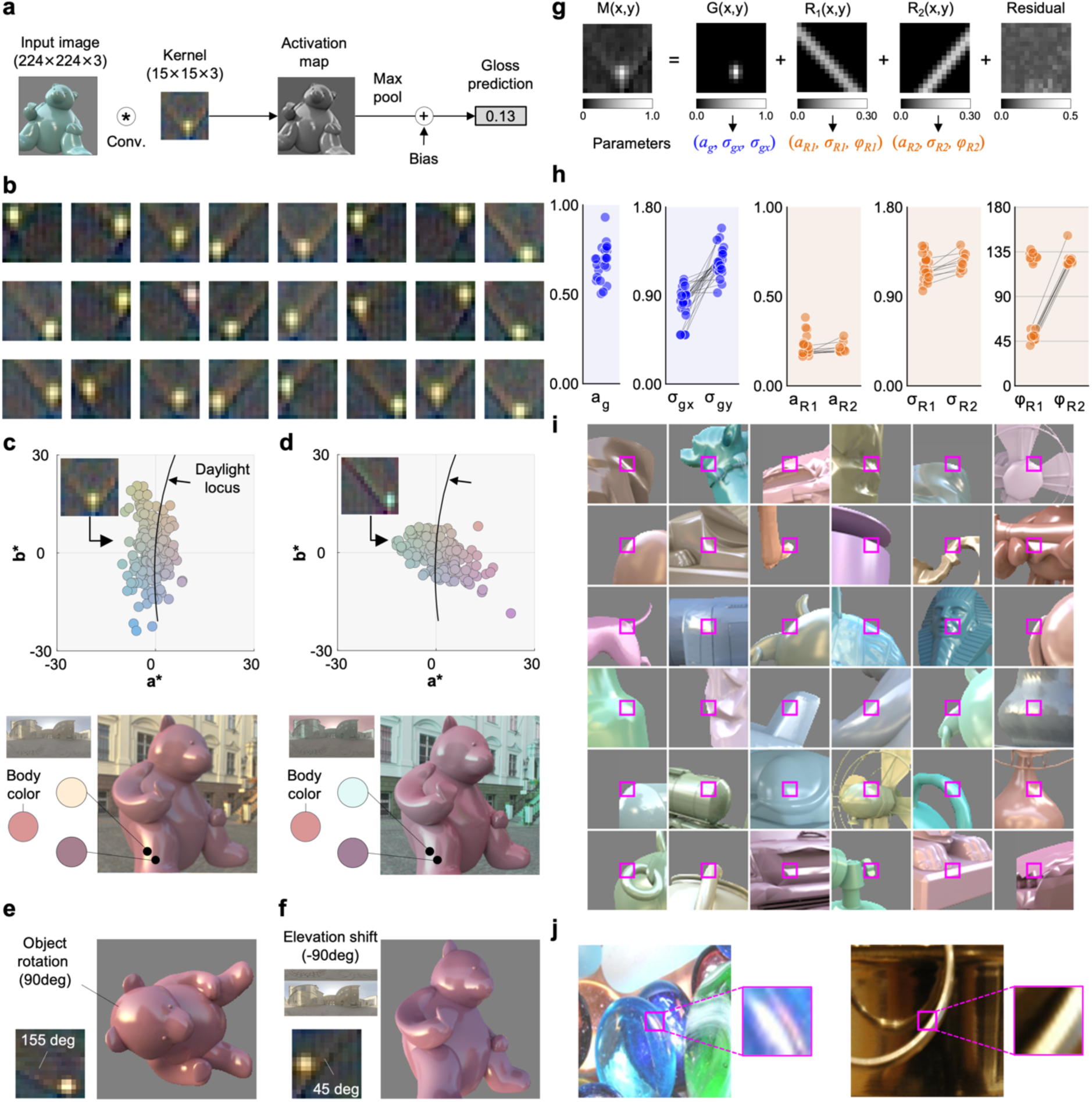
Analysis of single-kernel models trained on human gloss judgements. **a** Computational steps of the single-kernel model. The input image is convolved with a 15 × 15 × 3 kernel, the resulting activation map is max-pooled to a single value, and a bias term is applied to produce the gloss prediction. **b** Consistency check across 24 networks trained in cross-validation. Although they were trained on different (but overlapping) datasets with different random weight initializations, all networks developed a similar pattern of kernels, with a bright blob located at various positions, superimposed on straight elongated, diagonally-oriented brown-black-blue ridges. **c** Analysis of chromatic tuning of the kernel. The sRGB color of each pixel in an example kernel was converted to the *L*\**a*\**b*\* color coordinates, and their *a*\**b*\* distribution is shown. We found that kernel colors are closely aligned along the daylight locus, likely reflecting the typical color of specular highlights and their contrast with the surrounding bluish ambient light (from skylight reflections in shadows) seen in training images (see example image at the bottom). **d** Chromatic distribution of an emergent kernel from a human-like network trained on a new set of 3,888 images under 90° gamut-rotated environmental illuminations. The kernel clearly adapts to the chromatic statistics of specular highlights present in the training data. **e** Kernel that emerged from a human-like network trained on images in which the object was rotated by 90° relative to the original orientation. **f** Kernel that emerged from a human-like network trained on object images illuminated by light probes whose elevation was lowered by 90°. **g** Fitting a two-dimensional Gaussian filter and one or two Gaussian ridges to the kernel’s spatial intensity distribution. For simplicity, fitting was performed on the luminance image after mean subtraction. The bright blob region is well approximated by the 2D Gaussian filter. The diagonal stripe pattern is captured by a 1D Gaussian ridge, and the residual component is also shown. The difference in scale between the two components is indicated by the colorbars. **h** Distribution of the fitted parameters for the 2D Gaussian components and ridges across the 24 kernels shown in panel b. **i** Thirty-six image regions that most strongly activated the example kernel for images where both humans and the model judged the surfaces as highly glossy. Although the regions capture a variety of specular reflection geometries, diagonal highlights are comparatively more prevalent. **j** Examples of real material photographs, showing enlarged regions of oriented specular highlights that maximally activate our single kernel.

We found that the emergent kernels were highly consistent across the 24 networks (panel **b**). Their typical spatial structure consisted of a bright central blob surrounded by a darker region, which is well suited for detecting highlights based on their characteristic spatial intensity profiles. Notably, the bright region often exhibits a yellowish tint, while the surrounding area appears bluish. This mirrors the typical color relationship in natural scenes between direct illumination (such as sunlight) and indirect illumination (such as skylight), allowing the kernels to exploit hue differences for highlight detection. It is also notable that each blob is abutted by one or two brownish diagonal ridges—typically consisting of a brown upper region, a dark middle region and blueish lower region—presumably positioned to detect oriented, lower-contrast highlights. These ridges are bands of positive weights running at approximately 45° or 135°, which enable the kernel to respond strongly to local intensity changes at similar angles. As a result, the kernels can detect diagonal boundaries and curvatures typical of specular highlights on curved surfaces, as well as capture how highlights spread or extend in specific directions. Interestingly, while the “human-like” single-kernel networks (i.e., those trained on human labels) exhibited these distinctive diagonal ridges, the “ground truth” networks did not (see **SI** for example kernels). This suggests that the oriented ridges in the kernels truly capture something specific to human gloss perception—rather than being generically useful features for estimating reflectance. This may reflect the fact that the human visual system develops with a more varied visual diet and has to support many more visual tasks than simply estimating specular reflectance. Presumably, when asked to identify gloss, humans rely especially on features that distinguish specularity from other sources of image contrast.

Panel **c** shows the chromatic distribution of an example kernel from a human-like model in *L*\**a*\**b*\* color space. To obtain these coordinates, we converted the sRGB values of each kernel pixel to *L*\**a*\**b*\* using a D65 white point (X=95.0, Y=100, Z=108.9). The resulting distribution is tightly clustered along the CIE daylight locus, where natural illuminant colors typically fall [41]. As illustrated in Figure S3b, the environmental illuminations used in this study vary in gamut area but generally exhibit an elongated color distribution along the daylight locus, which directly influences the color of specular highlights as reflections of incident light. This suggests that the filters exploit this regularity to help extract specular reflections from object surfaces. To test this, we generated 3,888 new images with lighting environments in which pixel colors were rotated by +90° in the *a*b\** plane, while *L\** (lightness). All other rendering parameters matched the main dataset, and we assumed human gloss judgements would remain the same. As shown in **Figure 4d**, the chromatic distribution of the resulting kernel in the human-like model also rotated by roughly 90°, confirming that the kernel’s chromatic properties are shaped by the chromatic illumination statistics of the training environment.

To test whether certain geometric factors in the rendering process gave rise to the 45° and 135° ridge detectors, we manipulated object geometry and illumination to isolate their underlying contributions. When we rotated all objects by +90° relative to their original orientation (**panel e**), the angle of the ridge shifted to 155°, corresponding to a 20° change. In contrast, when we lowered the illumination elevation by 90°, moving the primary light source from the upper hemisphere toward lower regions, the kernel pattern exhibited little change (**panel f**). Therefore, the learned kernel ridge does not simply align with absolute lighting direction and overall is more affected by the projected surface geometry in the image set. See **SI** for extended analyses.

As shown in panel **g**, we found that the spatial properties of the bright blob and accompanying ridges can be represented by a two-dimensional (2D) Gaussian function, a type of receptive field observed as early as retinal ganglion cells [42] and one-dimensional (1D) oriented Gaussian ridges, respectively. The 2D Gaussian has five free parameters: amplitude (*a_g_*), x-center coordinate (*x_g0_*), y-center coordinate (*y*_g0_), standard deviations along the x-axis (*σ_g_*_x_) and y-axis (*σ_gy_*). Each 1D Gaussian ridge has five free parameters: amplitude (*a*_Rk_), x-center coordinate (*x*_Rk0_), y-center coordinate (*y*_Rk0_), standard deviations (*σ_Rk_*) and orientation (*φ*_Rk_), where k represents ridge 1 or 2. Although having both x-center and y-center is redundant (since the ridge functions are infinitely long), they were retained for consistency with the 2D Gaussian blob formulation. The correlation between the emergent CNN kernel and the fitted Gaussian blob and ridge model was 0.89 on average across 24 filters (SD = 0.017), implying that remaining local features in the residuals (rightmost image in panel **g**) contribute minimally to the overall structure.

Panel **h** shows the fitted parameters for each function across the 24 kernels displayed in panel b, excluding *x_Rk0_* and *y_Rk0_*. The 2D Gaussian blobs exhibited high amplitude values (0.5–1.0) and standard deviations indicating a slight vertical elongation. Among the 24 kernels, 9 were fitted with two ridges. The amplitudes (*a_Rk_*) of these ridges were consistently lower than those of the Gaussian blobs, typically ranging from 0.2 to 0.3, and *σ_Rk_* values were similar between ridges. The orientation parameter (*φ_Rk_*) clustered clearly around 45° and 135°. When two ridges were present, they were separated by 90°, indicating a systematic orientation preference in the filter.

We examined the local image features that most strongly activate the “blob-and-ridge” kernel (panel i). Each image shown was judged as highly glossy by both the model and human observers, and the magenta square marks the local patch that maximally activates the kernel. These examples reveal that strong activations typically occur for regions containing an elongated, collinear highlight “ridge,” most often oriented diagonally (∼45° or 135°). Unsurprisingly, the kernel also responds to more circular highlight structures (“blobs”) (e.g., fifth row, third column). Another notable observation is that the filter is sensitive to cases where the highlight terminates abruptly at an object boundary or sharp surface edges. Such configurations are rarely produced by diffuse (Lambertian) shading in natural lighting environments, which instead generates smoother intensity gradients. At the same time, these examples also illustrate that the kernel can respond to a variety of specular highlight patterns.

We emphasize that the strong orientation selectivity of the kernel is unlikely to be merely an artifact of our training image set. Oriented specular highlights are frequently observed in everyday materials that contain roughly cylindrical structures, in which the highlight is necessarily elongated, as demonstrated by examples taken from the Flickr Material Database (FMD) [43] (see panel **j**). This pattern is thus characteristic of real-world surfaces, not just synthetic images. Further validation of the single-kernel model’s performance on real-world photographs is presented below (see **Figure 8**).

In summary, the kernels observed within the tiniest CNN are composite filters that encode multiple fundamental visual features, including luminance and chromatic contrast, orientation, and shape. This allows them to capture both the presence and geometric structure of specular highlights, supporting robust extraction of highlights across a range of different geometries. Although such a simple filter was far from the best of the models we tested for predicting human gloss judgments, it is striking that its correlation coefficient reaches 75.3 percent of the upper-limit set by inter-observer consistency, yet is highly interpretable, yielding insights into key features that observers likely rely on to make their gloss matches.

### Analysis of three-layer models

**Figure 5a** illustrates the computation flow of the example three-layer model with 64 kernels. The model processes the input image through multiple layers of convolution, pooling, normalization, and non-linear activation, ultimately using linear regression to predict the gloss value. The leftmost images show nine example kernels in the first convolutional layer, which show Gaussian blobs with different spatial scales, contrasts, color tunings, and polarities.

**Figure 5.**
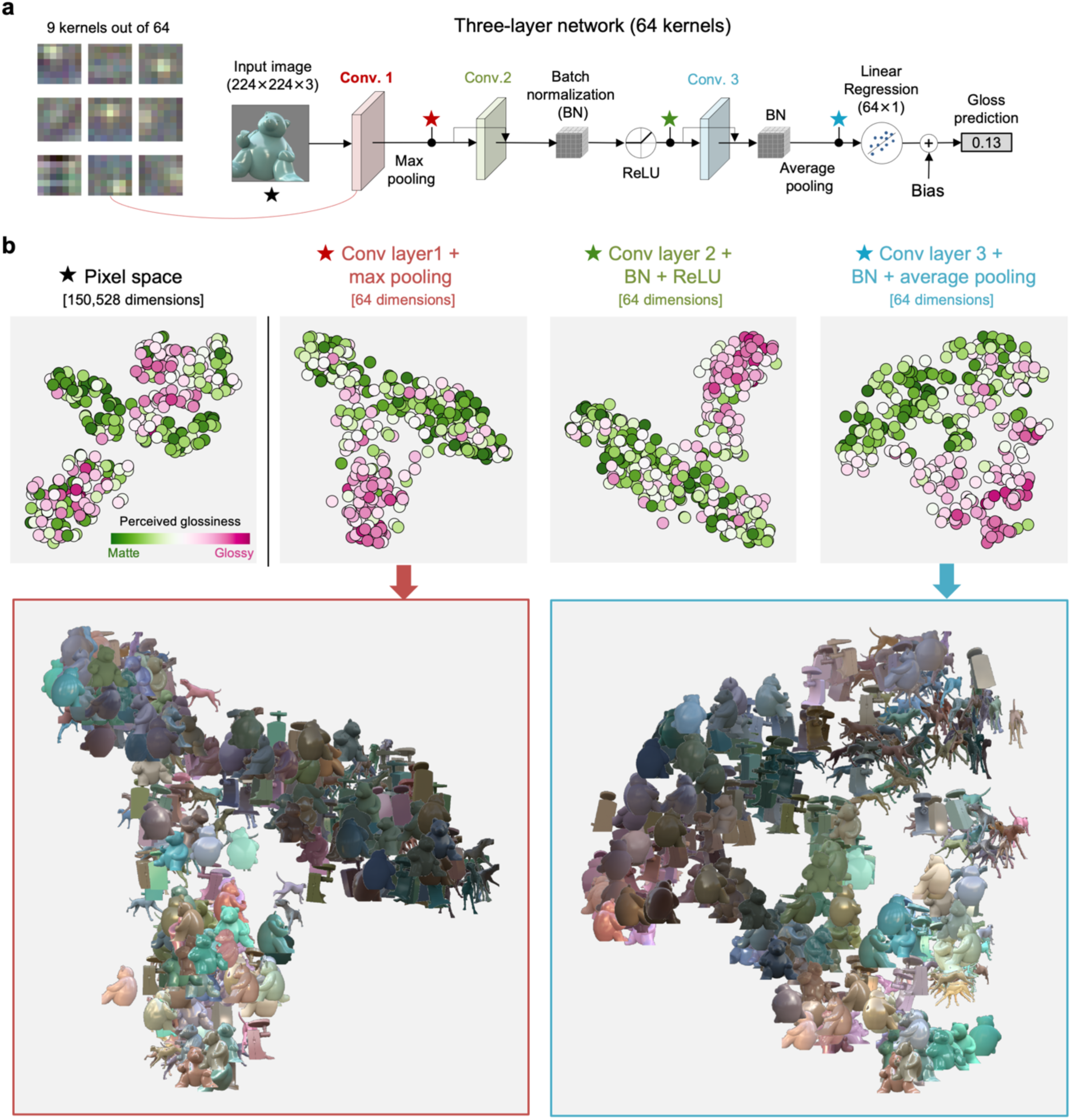
Analysis of a three-layer model with 64 kernels trained on images with human labels. **a** In the first convolutional layer, the input image was convolved by a set of 64 kernels, 9 of which (7 by 7 pixels) we visualize on the left side. In the second and third convolutional layer, activation maps from the first convolutional layer were further convolved by the two sets of kernels subsequently and average pooling was taken for each of 64 activation maps from the third convolutional layer. Between convolutional layers 2 and 3, batch normalization and/or ReLU are applied. There were skip connections between layers to bypass the information if beneficial. Then the pooled features were used as input for a linear regression model, which predicted gloss level. **b** Example of internal representation in a three-layer network model. We fed the network images it had not seen during training and extracted activation maps from each convolutional layer. The maps were aggregated by max pooling (layers 1 and 2) or average pooling (final layer) over the entire map, then used as inputs for t-distributed stochastic neighbor embedding (t-SNE) to project onto a two-dimensional plane. The bottom two figures show representations from the first and third convolutional layers. For comparison, we also performed t-SNE analysis on the same images in the pixel space (leftmost plot), where matte and glossy objects are not separated and objects are primarily clustered by shape (not shown here).

We examined this three-layer network, the best of all the models we tested, to see how glossiness is coded within the network and how the representation progressively changes over layers, using 324 images that were not included in the training dataset for this network. We first extracted activation maps after each convolutional layer (marked by stars) and aggregated the signals over the map using max pooling for the first layer and average pooling for the third layer. This resulted in 64 scalar values per layer, which were plotted on a 2-dimensional plane using t-distributed stochastic neighbor embedding (t-SNE; panel **b**). For comparison, we performed the same analysis using the raw pixel values of the input image.

Results revealed that in pixel space, objects are primarily clustered by their geometry (the plot does not label specific shapes). In the first layer, object images are already roughly grouped into glossy and matte categories, with some glossy objects misclassified as matte and vice versa. The second and third layers only slightly refine this representation. However, the overall representation does not change drastically from the first layer. To quantify this progression, we trained a linear regressor with 10-fold cross-validation to predict human gloss judgments from each stage’s readout. Pearson’s correlations (mean ± SD), averaged over 10 folds and 24 networks, were 0.13 ± 0.13 for pixel space, 0.64 ± 0.074 for layer 1, 0.67 ± 0.076 for layer 2, and 0.79 ± 0.0586 for layer 3. A one-way repeated-measures ANOVA revealed a main effect of layer (*F*(3, 69) = 307.1, *p* < 0.001, Cohen’s *f* = 3.65). Bonferroni-corrected pairwise comparisons (α′ = 0.0083) confirmed the following significant differences: pixel space < layer 1, layer 2, and layer 3; layer 1 < layer 3; and layer 2 < layer 3. This suggests that the first convolutional layer most strongly contributes to gloss judgments, with subsequent layers making small but significant adjustments to improve prediction, as well as arranging objects by their shape characteristics (note the bulbous-spiky organization [44, 45] in the layer 3 representation)

Taken together with the analysis of the single-kernel network, these observations suggest our gloss judgments can be predicted well by a combination of well-known, low-level filtering mechanisms within the visual system.

### Evaluation of model generalization across supplementary image sets

We conducted three additional generalization tests using (i) object images with manipulated specular highlights, (ii) an additional set of 42,120 rendered images, and (iii) real-world photographs, as detailed below.

### Gloss illusion caused by manipulated specular highlights

Surprisingly, given the simplicity of the internal representations, we also found that the human-like networks predicted a number of known gloss effects [8, 46, 47] as shown in **Figure 6**. For instance, the networks accurately anticipated the decrease in perceived glossiness when the specular highlight is rotated (panel **a**), moved horizontally (panel **b**), when surface roughness increases (panel **c**), and when the underlying specular reflectance weakens (panel **d**). The decrease was steeper for the three-layer models compared to the one-layer model. Notably, these modified images were not included in the training dataset, demonstrating the models’ ability to generalize beyond the training range. It is straightforward that perceived gloss decreases when surface roughness or specular contrast decreases, as this correlates with a decrease in local intensity of the specular component. It is somewhat surprising, however, that even a single-kernel model can predict the decrease in gloss level for rotation and translations. The spatial pattern of the receptive field suggests the filter utilizes the local spatial intensity profile of the specular highlight region. For a typical glossy object, this local intensity profile likely arises from the combined contribution of both specular highlights and diffuse components when they are correctly aligned [48]. When only the specular component shifts from the original position independently of the diffuse component, this geometrical regularity collapses, leading to a decrease in the filter output. However, we recognize this is not always the case. Out of 3,888 test images, 790 images (about 20%) actually showed an overall increase in gloss levels due to highlight rotation. Nevertheless, it is intriguing that simple models can detect seemingly complex inconsistencies of highlight position. It suggests that while there is surely some degree of coupling between mid-level representations of shape, material and lighting [5, 9, 12, 13, 49, 50], a significant portion of gloss perception may actually be accounted for by more basic mechanisms.

**Figure 6.**
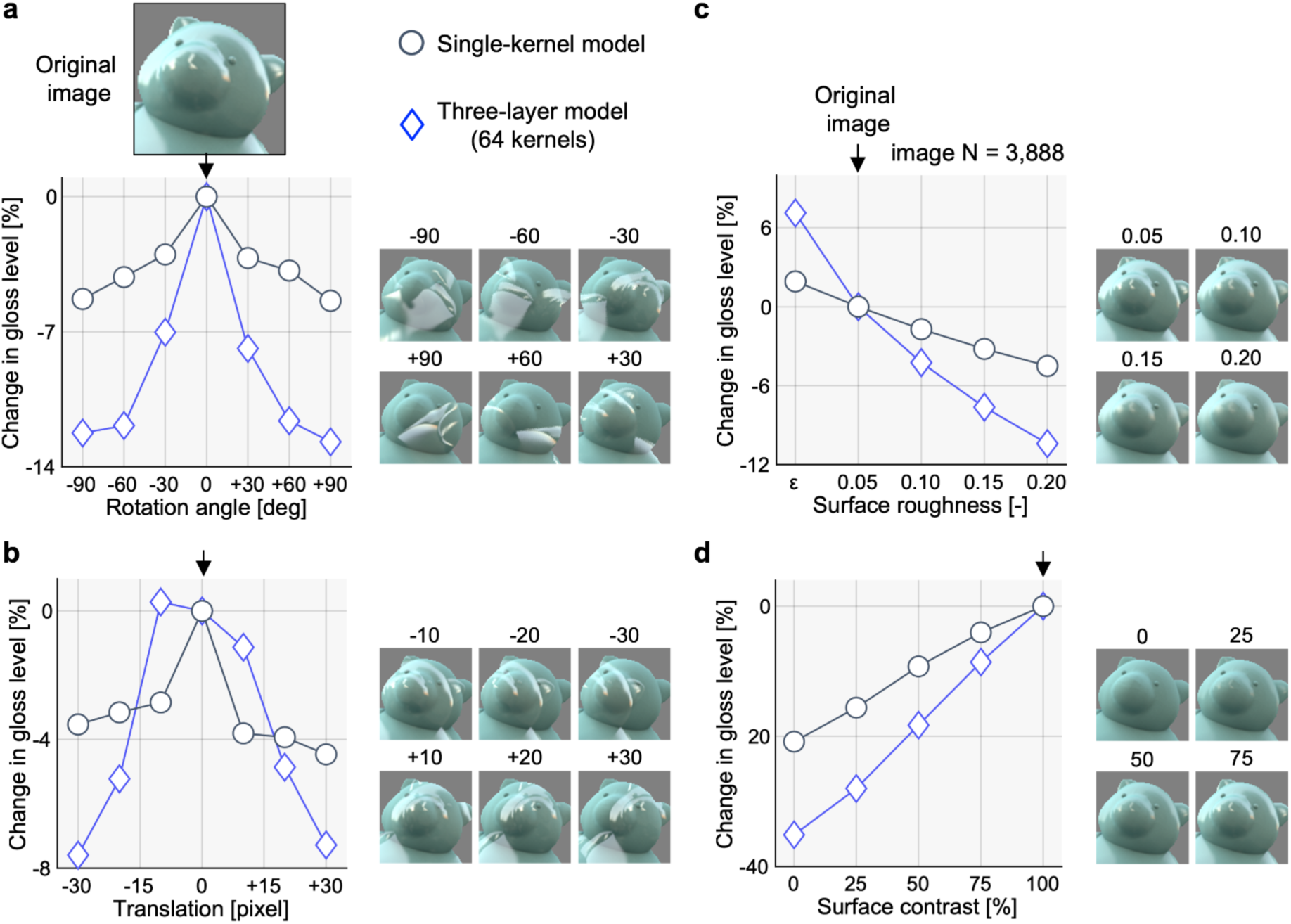
Evaluation of single-kernel and three-layer network models for three known perceptual effects. The dark blue circles and blue diamonds show model responses for the one-layer model and three-layer models, respectively. **a** Effect of rotating specular structure. The top image shows the original image where no manipulation was applied. The rotation 0 indicates the original image. **b** Effect of horizontally translating specular reflection. The translation 0 indicates the original image. **c** Effect of changing surface roughness level. The surface roughness 0.05 indicates the original image. **d** Effect of changing surface contrast level. The surface roughness 100% indicates the original image. Contrast was manipulated by changing Pellacini’s c.

### Additional rendered images

Serrano et al. [51] generated 42,120 images from 9 lighting environments (**Figure 7a**), 9 object geometries (panel **b**), and 520 bidirectional reflectance distribution functions. For each image, they collected ratings of various perceptual attributes including glossiness through crowdsourcing, providing an excellent validation set for our models. We tested our single-kernel model and three-layer model with 64 kernels, both trained on all 3,888 images. To prepare the input, we used object masks to exclude non-object regions, filling them with a mid-gray color (sRGB = [127, 127, 127]), and then fed the resulting images into our models to generate gloss predictions.

**Figure 7.**
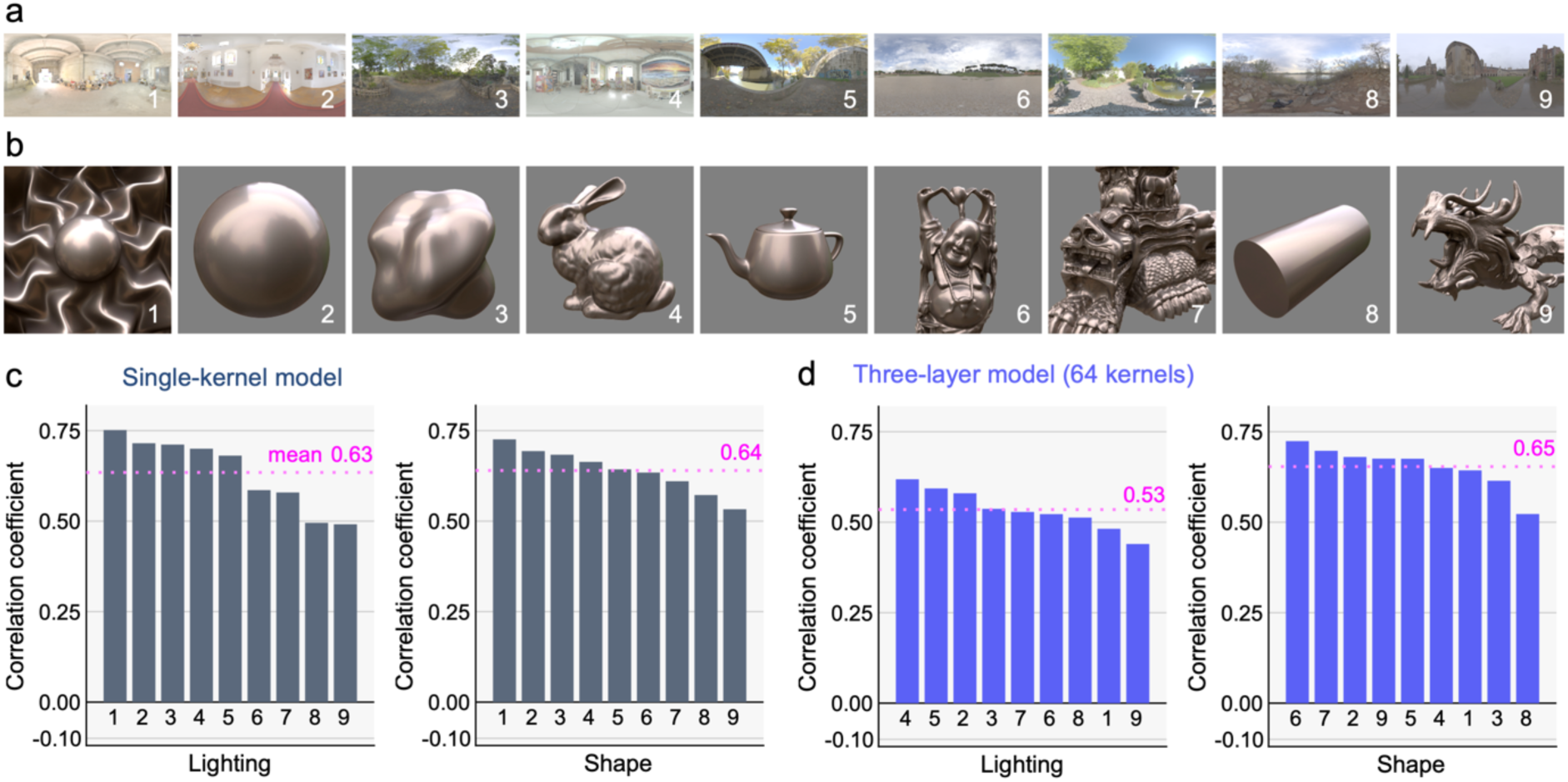
Results of a generalization test using Serrano dataset. **a, b** Nine lighting environments and shapes used to generate rendered images. **c** Pearson’s correlation coefficient between the model response of the single-kernel model trained by all 3,888 images and perceptual rating measured in crowd-source experiment [50] for each lighting and shape. Each bar shows the correlation across 4,680 images. **d** The case for a three-layer model trained by all 3,888 images, which shows slightly lower generalization ability compared to the single-kernel model.

Panel **c** shows Pearson’s correlation between the single-kernel model predictions and human gloss ratings. The single-kernel model, despite its minimal architecture, achieves a mean correlation of 0.63 (SD = 0.099) and 0.64 (SD = 0.061) across lighting environments and shapes, respectively, performing comparably to our own dataset (mean *r* = 0.65). There is a general trend that geometries with smoother surfaces are better predicted by the model, although there are some exceptions, such as the bunny (geometry 4).In contrast, the three-layer model shown in panel **d**, which outperforms the single-kernel model on our own data (mean *r* = 0.82), shows a correlation of 0.53 (SD = 0.056) and 0.65 (SD = 0.058). Similarly, the deep ResNet-52 gloss model by Serrano et al., when applied to our 3,888 images, shows a weak correlation of 0.29 (not shown), highlighting a trade-off between model complexity and generalization. Clearly human perception does not suffer from such generalisation limits, yet this likely relates to our vastly larger visual diets.

### Real-world photographs

We further evaluated generalization to real-world photographs of everyday materials using 183 images compiled from multiple sources, including the Flickr Material Database (FMD) [42], visuo-haptic studies by Baumgartner et al. [52, 53], an fMRI study by Jacobs et al. [54], and a perceptual material classification study by Wiebel et al. [55]. The dataset covers a diverse range of material categories, such as metal, glass, plastic, fur and leather, fluids, and fabric. All images had been consistently classified as either matte or glossy by all eight observers in a previous study [11].

**Figure 8a** shows the predictions of the single-kernel model and the three-layer model, plotted on the x-axis and y-axis, respectively. The classification accuracy, based on the threshold indicated by the dashed lines in the plot, was 91.9% for the single-kernel model (*d’* = 2.79; a bias-free measure of discrimination sensitivity) and 71.9% for the three-layer model (*d’* = 1.08). This again demonstrates the superior generalization performance of the simpler model. Notably, the misclassified images lie close to the category boundary, with a mean distance of 0.0079 (SD = 0.0081), which corresponds to approximately 7 % of the model’s prediction range.

**Figure 8.**
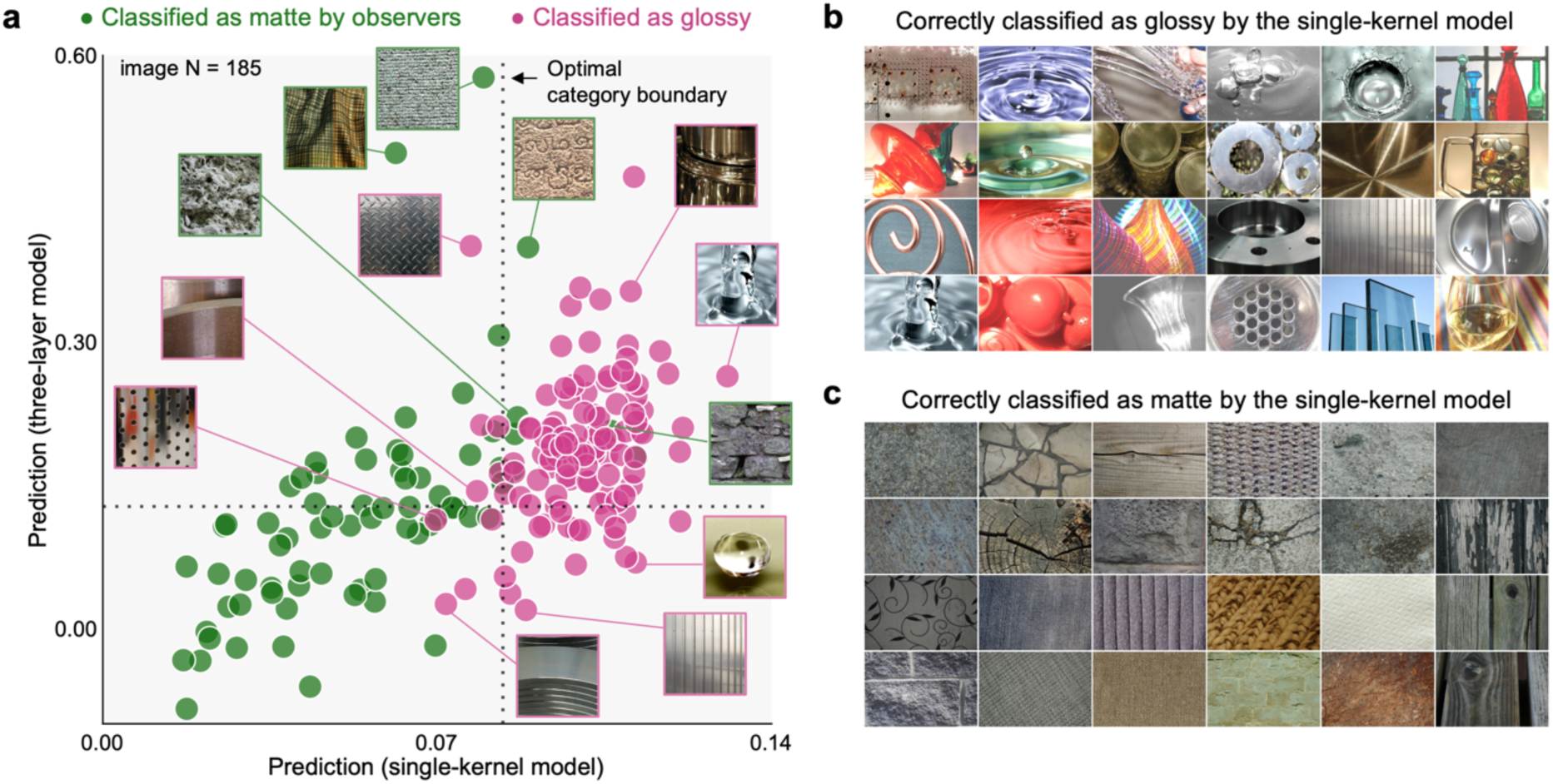
Generalization performance on real-world material photographs. **a** A scatter plot comparing model responses for each image labeled as either matte or glossy based on behavioral data from a previous study [11]. The x-axis represents the single-kernel model, and the y-axis represents the three-layer model. Green and pink circles show images consistently rated as matte and glossy, respectively, by all eight observers. The dotted lines indicate the optimal category boundaries that achieve the highest classification accuracies: 91.9% (*d’* = 2.79) for the single-kernel model and 71.9% (*d’* = 1.08) for the three-layer model. **b** Example images correctly classified as glossy by the single-kernel model. **c** Example images correctly classified as matte.

This result is intriguing in that even a simple filter-based model can capture the diverse patterns of specular reflection found across different real-world materials (panel **b**) with high accuracy, while remaining robust against being misled by the texture patterns present in many matte images (panel **c**). Interestingly, the three-layer model, which performed well on our own dataset, showed reduced accuracy in this test. This suggests that even relatively shallow models, such as those with only three layers, can be prone to overfitting, pointing to the often-overlooked value of using simpler computational strategies.

Together with the results on manipulated specular highlights and Serrano’s 42,120 images, this validation shows that the single-kernel model achieves an effective balance between simplicity and performance, making it a strong candidate for gloss computation across diverse image sets.

## Discussion

This study is inspired by a fundamental question in behavioral neuroscience: what neural computations enable material perception from highly variable sensory inputs across different contexts? We address this by leveraging CNNs to replicate human-like judgments of object gloss from images and explore the internal computational strategies of these networks, in a data-driven, hypothesis-neutral approach that contrasts with traditional cue-based methods. The motivation is that tasks such as material perception might be too complex for intuitive identification of underlying mechanisms, whereas CNNs excel at finding statistical regularities in training datasets. We used this tool to guide our search for biologically plausible computations that underlie human gloss perception.

This approach identified a clear contrast between the computational demands required to develop human-like CNNs and physical ground-truth CNNs, suggesting that human gloss judgments from object images likely do not involve sophisticated inverse optics computations. This notion has been suggested previously [1], but what is new here is the identification of specific filters and the depth of computation required to predict human gloss perception and physical reflectance parameters. In particular, the effectiveness of a filter with a bright blob and ridges is intriguing as it appears to capture a wide range of specular structures despite its simplicity. This finding directly supporting the idea that gloss perception relies, to a nontrivial extent, on relatively simple image features, as suggested by Motoyoshi et al. [6]. Similarly, previous studies suggested that the visual system may use the steepness of the intensity gradient to separate shading into shape and glossiness components [56, 57]. Much as cone photoreceptors provide foundational signals for color perception and other visual functions, bright blob and ridge detectors may serve as a key component of gloss perception, and perhaps other distal perceptual attributes too [58–60]. In light of this, one hypothesis is that low-level computations supply a compact set of general-purpose image features that can be efficiently recombined through flexible, task-dependent selection mechanisms—gloss perception in the present case—underpinning the remarkable versatility of human material perception.

While we certainly are not suggesting that the human gloss perception is based *exclusively* on low-level filters, it is intriguing that when a large population is asked to judge gloss under diverse conditions, they resort to strategies that can be well approximated by such measurements. We would caution against interpreting our findings as suggesting the human visual system contains ‘gloss detector’ filters resembling the kernels that emerged in our models. Representations in the human visual system have to underpin far more tasks than just gloss perception. Instead, we see the emergent kernels as a data-driven means to identify the relevant *image information* that observers draw on to perform the gloss match task while noting that multi-kernel models (3-layer, 64 kernels) better predict human judgments overall. Yet the combination of controllable computer-generated training sets, large-scale human perceptual data, and deep learning with small, interpretable models seems to be a promising avenue for unravelling the computations underlying other ‘mid-level’ visual inferences.

The diagnosticity of diagonal specular reflections is also consistent with an ecological perspective, given the distribution of orientation signals in natural images [61]. Since vertical and horizontal edges are pervasive in the environment, these axis-aligned contrasts are so common that their presence alone provides little information about gloss. In contrast, diagonal signals are rarer, so their presence in the image is a stronger cue that the cause is a specular reflection. Interestingly, the geometry 1 used in Serrano’s dataset (**Figure 7**) is known to produce a particularly strong gloss impression [62], and its spatial pattern resembles the feature captured by our single kernel. Consistent with this, our model predicted human gloss judgments most accurately for this geometry. Finally, although the present study focused on isotropic materials, many natural and artificial materials—such as brushed metals, silk fabrics, and polished wood—are anisotropic and often produce directionally elongated specular highlights, suggesting that the mechanisms captured by our kernel model may be applicable to these material types as well.

One key aspect of our approach is the use of large-N psychophysical experiments. Since numerous distal factors such as shape, lighting, and reflectance properties are involved in generating a single object image, covering a wide range of stimulus space—ideally, comparable to the diversity of natural images—is critical to capture visual behaviors with sufficient resolution. Conversely, when the analysis was limited to a small number of shapes or lighting environments, some cases happened to show relatively high correlations between perceptual gloss judgments and the physical ground truth (**Figures S4 and S5**). This correlation is not inherent to our perceptual judgment but is influenced by the stimulus set. Moreover, it is worth noting that behavioural error pattern we observed shows a strong asymmetry: as illustrated in **Figure 2e**, there are far fewer cases of “false positives” (objects that are not physically shiny but appear so) than “misses” (high-reflectance objects that do not look glossy). This likely arises from basic optical constraints—it is difficult to produce salient specular highlights on low-reflectance objects, whereas it is relatively easy to eliminate highlights on high-reflectance ones; for example, by placing a glossy object in a diffuse lighting environment or rotating a planar glossy object so that it reflects the ground rather than the sun can make a shiny object appear matte. Recent studies have made significant efforts to collect extensive human judgment data across a large number of images [63, 64]. To understand visual mechanisms, we must account for both successes and errors in perception, and large-scale measurement helps capture these diagnostic behavioral patterns. Although careful data curation and validation are essential to this approach, it offers a powerful data-driven framework for exploring the complex mechanisms underlying human perception and behavior. Combining our approach with unsupervised learning has also shown some potential for reducing the cost of collecting labelled images [65–67].

There is a general concern about the lack of calibration and control over viewing conditions in online experiments. Our stimuli were presented assuming an sRGB display profile, but individual observers’ monitors may deviate from this assumption (e.g., in chromaticity or gamma of each RGB phosphor). Importantly, however, our glossiness was measured using an asymmetric matching task—a relative judgment in which both test and reference images were viewed under the same display conditions. Thus, even if an observer’s monitor had an atypical gamma or contrast profile, it would have affected both images similarly and is unlikely to have systematically biased their settings. Moreover, as shown in Figure S6 (laboratory experiment), we observed a high degree of consistency between online and offline data, suggesting that variations in viewing conditions did not substantially influence observers’ gloss matches.

A related point is that neither online nor laboratory-based experiments employed a high-dynamic-range (HDR) scene. To ensure compatibility with standard monitors, we deliberately avoided extremely high dynamic range lighting environments and viewpoints producing strong specular highlights. The maximum specular reflectance was capped at 0.0999 (9.99%), which still yielded highly glossy appearances. Using a higher value could allow a wider range of glossiness to be explored, as demonstrated in recent HDR display work [68]. This constraint is shared by most material perception studies and shows an avenue for future investigation.

Our model validation results highlight the generalization challenges inherent to complex models. While deep learning has addressed many challenges in visual perception, it remains uncertain whether such models capture the mechanisms underlying human vision [69]. A key strength of our approach is its capacity to systematically explore specific filter designs and their combinations, enabling a more comprehensive search through the space of low-level computations. Our results may provide insight into long-standing questions in vision science, specifically whether complex perceptual systems like material perception originate from low-level visual processes [70].

One outcome of this study is the creation of a publicly available database containing 3,888 images. While many databases exist for labeled objects, those focusing on material properties are less common. Each image is annotated with full stimulus parameters, including physical ground-truth labels and perceptual labels from at least three different observers. We expect this dataset to be useful in multiple disciplines such as vision science, cognitive neuroscience, and computer vision. In addition, the availability of perceptual labels is useful for industrial applications, such as predicting the perceived glossiness of car paint.

To conclude, our study has provided a novel data-driven perspective on the computational mechanisms in human gloss judgements using large-scale perceptual measurement and CNN-based modeling. We found a relatively simple and biologically plausible linear model that predicts the idiosyncrasies of human gloss constancy, but also several established perceptual gloss effects beyond the training range, as well as gloss perception for photographs of real-world objects.

## Materials and Methods

### Stimuli

The detailed procedure for stimulus generation is explained in our previous study [30]. All reference and test images were generated using the physically-based rendering software Mitsuba v 0.6 [71]. Images were generated in XYZ 1931 format. For the online experiment, the image was converted sRGB and gamma correction was applied. All images contained a single object at the center of the scene, to which we applied diffuse and specular reflectance using the Ward reflectance model [37, 38] with a fixed surface roughness level of 0.05. The specular reflectance value was randomly sampled from 0.0029 (weakly glossy) to 0.0999 (highly glossy). The thirty-six objects consisted of 3D models of varying complexity, including both 3D scans of real objects and artist-created models, spanning natural and man-made artefacts and enabling us to capture how different geometries influence highlights and other indicators of gloss. We applied image-based environmental illumination [72] to the scene. Each lighting environment had been photographically captured from a specific real-world location, with each pixel representing the colour and intensity of light arriving from a particular direction at a single point in space—thereby capturing illumination from all directions. We used thirty-six lighting environments, collected from the Debevec database [72], Southampton-York Natural Scenes [73], and Freebies (https://hdrmaps.com/freebies/; accessed May 2021). Further details are provided in the **Supplementary Information** (Figure S3). Image-based environmental illumination was set at infinity, with the camera positioned at the object’s height and facing it directly. Interreflections were included in the rendering. The object was randomly rotated around the Z-axis, while rotations around the X- and Y-axes were limited to ±15°, maintaining an upright orientation but allowing for some variation (see **Figure 1a** for the XYZ rotation axes). We deliberately avoided scenes with extremely high dynamic range lighting, allowing the images to be displayed on standard monitors without substantial tone compression. The asymmetric gloss matching task required a matching object whose physical specular reflectance parameter could be continuously adjusted by observers. For this, we used Pellacini’s c [38], a perceptually-linear characterization of Ward specular reflectance associated with the object to render a set of images, instead of making changes in image space. A total of 3,888 test images were generated by combining 36 lighting environments, 36 object shapes, and 3 random viewpoints. For each test image, a random body color was used, with fixed lightness at 50, and random chroma and hue between 8-26 and 0°-360° in the L*a*b* perceptually-uniform color space [74], respectively. A full list of object shapes and environmental illuminations is shown in **Figure S2 and S3**.

To generate the textured object images used for validation, we selected 12 object geometries from the original set of 36 that had smooth, continuous surfaces suitable for texture mapping. We then created 1,296 images by applying achromatic blob textures with either the same luminance as the body color or 1.5 times higher (**Figure S8a**).

All images were generated at 512×512 pixels in XYZ format. After converting the raw XYZ image to a linear monitor RGB format, each image was normalized using the 97th percentile pixel value. These scaled linear RGB images were then gamma-corrected using the standard sRGB formula. The spatial resolution of 512×512 pixels was used for the psychophysical experiment, but reduced to 224×224 pixels for the network’s inputs to shorten the training time.

For the online experiment, the total of 3,888 images were divided into 54 independent sets. Twelve common images were added to each set to evaluate the consistency of observer responses. At least three observers were recruited for each set. For two of these sets, we recruited 21 and 23 online observers, respectively, and compared their responses with those of 20 observers from the lab-based experiment to validate the online data (see **SI**).

### Experiment

The main experiment was conducted online using PsychoPy (version 2022.1.4) [75] and hosted on an online platform Pavlovia. Observers were recruited via Prolific and compensated 12 euros per hour. Before participating, they downloaded and read an information sheet explaining the procedure and signed a digital consent form. A validation experiment was conducted in the laboratory (detailed in SI, Figure S4).

### Observers and Ethics

A total of 467 observers participated in the online experiment, with data from 295 analyzed (see exclusion criteria in a later section). Observers had normal or corrected-to-normal visual acuity and normal color vision (self-report). Their age ranged from 19 to 65 (mean 41.4; S.D. 12.4). The observer group comprised 159 females and 136 males. Informed consent forms were in English, so only English-speaking observers were recruited for the online experiment. No language criteria were applied for the laboratory experiment. No observer was informed of the experiment’s hypothesis. The study was approved by the local ethics committee at Justus Liebig University Giessen, following the Helsinki Declaration (sixth revision, 2008).

### Procedure and task

Observers were instructed to turn off the room lights and sit at arm’s length from the display during the experiment. Observers viewed the screen binocularly. Before the experiment, observers completed a size calibration using a bank card [76] to ensure the stimulus image appeared at the correct size. In each trial, a test image was shown on the left and a reference image on the right. The task of observers was to move a slider under the reference image to match its perceived glossiness with the test image. The reference image stayed the same in all parameters except Pellacini’s c, while the test image varied between trials in terms of shape, color, viewpoint and lighting environment (asymmetric matching [77,78]). After completing the size calibration, the observers received on-screen instructions: “Adjust the gloss level of the right ‘reference’ object using the slider until it matches the gloss level of the left ‘target’ object.” The slider’s starting position was randomized for each trial, excluding edge values, which were reserved for catch trials used to detect autoresponders. The first three trials were practice rounds to help observers get used to the task. After practice, they adapted for one minute to a uniform white screen (sRGB = [127, 127, 127]). Each observer was randomly assigned to one of 54 image sets, each containing a set comprising 84 images (72 unique to the set and 12 common images). Each session consisted of 84 trials, and all observers completed 2 sessions, totaling 168 trials. To detect autoresponders, each session included a catch trial in which no stimuli were shown; instead, a text screen instructed observers to move the slider to the left edge before proceeding to the next trial. The order of stimulus presentation was randomized. Median time across 295 observers to complete the experiment was 13.7 mins (4.89 sec per trial).

### Exclusion criteria

For quality control, we set two exclusion criteria for online experimental data. First, observers who failed either of the catch trials were excluded. Second, observers who completed each trial too quickly were excluded; this was indicated by a median response time shorter than the median minus two standard deviations of laboratory-based experiments. The rationale is that a matching task requires comparison and adjustment; decisions at implausibly short latencies are a strong indicator of inattentive or automated responding. We continued our experiments until at least 3 observers passed these criteria for each image set. This exclusion process resulted in 295 out of 467 observers (63.2%) being included in the final analysis. In contrast to the online experiment, no data were excluded from the laboratory validation experiment.

### Computational Modeling Fitting of single-kernel models

To quantitatively describe the spatial structure of the filters developed in network models, we modeled them as the sum of a two-dimensional (2-D) Gaussian *G*(*x*,*y*) and multiple oriented ridge functions *R_k_*(*x*, *y*) as defined in equation (1):

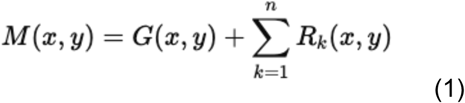

The 2-D axis-aligned Gaussian function was defined as equation (2):

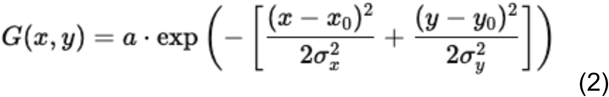

where *a* is amplitude, (*x*_0_,*y*_0_) is the center, and *σ*_x_, *σ*_y_ are the standard deviations along each axis.

Each oriented ridge was defined as equation (3):

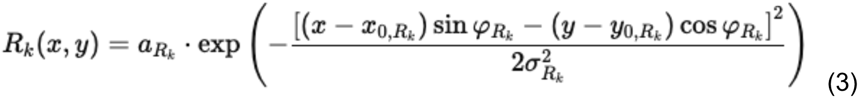

where *a_Rk_* is amplitude, (*x*_0,Rk_, *y*_0,Rk_) is the ridge center, *σ_Rk_* is the width perpendicular to the ridge, and *φ_Rk_* is the orientation. The index *k* takes values 1 and 2.

We converted each kernel from sRGB to XYZ and used the Y (luminance) channel for fitting to simplify the analysis. Optimization was performed using MATLAB’s lsqcurvefit function. The functions were fitted sequentially: first the primary ridge, then the secondary ridge, and finally the 2D Gaussian. If a second ridge component was not identified through the optimization, only a single ridge was fitted to the kernel.

### Luminance statistics models

The Y (luminance) channel from raw XYZ images was extracted. We implemented nine models to determine glossiness using the mean, median, standard deviation (std.), skewness, kurtosis, first quartile (Q1), third quartile (Q3), and the minimum and maximum pixel luminance values within the object region of each test image, excluding the surrounding region from these calculations.

### Specular reflection models

We computed three metrics—coverage, sharpness, and contrast—derived from the specular reflection component of each image, inspired by an influential prior study [12]. Each model involved free parameters, which were optimized to align most closely with human gloss judgments. For each test image, we rendered a version of the object containing only the specular component by setting the diffuse reflectance to zero, and converted the resulting image to a luminance image. To extract direct specular reflections and exclude secondary and higher-order interreflections, we selected only those pixels whose intensities exceeded a certain percentage (*k*%) of the maximum intensity in the specular image. The value of *k* was selected from the set {0, 1, 3, 5, 10, 20, 40}. Based on this thresholded highlight image, we computed the three metrics. Coverage was defined as the proportion of the object’s region occupied by highlights. Sharpness was calculated using spatial convolution to detect regions with rapid luminance changes [79]. For contrast, rather than using the raw highlight image, we applied a Gaussian band-pass filter to decompose the image into seven spatial frequency sub-bands, with cutoff ranges of 1.5–3.0, 3.0–6.0, 6.0–12.0, 12.0–24.0, 24.0–48.0, 48.0–96.0, and 96.0–192.0 cycles per image. This approach follows previous findings suggesting that certain frequency channels contribute more significantly to gloss perception [80]. We then computed the root-mean-square contrast, equivalent to the standard deviation of pixel intensities, for each sub-band image, as well as for an aggregated image across all frequency bands. Unlike coverage and sharpness, which each depend on a single parameter, the sub-band contrast metric includes two free parameters: the pixel intensity threshold *k* and the spatial frequency band. These parameters were optimized separately for each metric to determine the values that best correlated with human gloss judgments.

### Manipulation of the pattern of specular reflection

First, for each of the 3,888 test images, the diffuse and specular components were rendered separately. We then manipulated the specular reflection in three different ways (rotation, translation and roughness) before combining it with the diffuse component. The strength of the specular reflectance was set to a maximum value for all images (0.0999 in the Ward reflectance model, equivalent to 0.1487 in Pellacini’s c) to clearly observe the effect of highlight manipulation.

For rotated specular reflection, we rotated the specular component from −90° to +90° in 30° steps, using the image center as the rotation axis. For translation, we shifted the specular component horizontally from −30 to 30 pixels in 10-pixel steps relative to the image size of 224×224 pixels. After these manipulations, a mask was applied to remove any specular reflection extending beyond the object region. For surface roughness, we rendered the specular component with varying roughness values of ε (0.001), 0.05, 0.10, 0.15, and 0.20. For surface contrast, we adjusted the strength of the specular component between 0, 0.0372, 0.0743, 0.1115, and 0.1487 in Pellacini’s c unit. Then, we combined each manipulated specular image with the corresponding diffuse image.

All manipulated and original images were input to network models, and the proportional change in predicted gloss level relative to the non-manipulated image was computed.

### Network architectures

We chose a ResNet architecture for its effectiveness in object recognition tasks [81] and its modular design, which enables systematic adjustment of architectural complexity.

The one-layer model consists of a single convolutional layer, followed by max pooling and a regression layer to predict continuous gloss values. It does not include non-linear operations such as rectified linear unit (ReLU) or batch normalization. The three-layer model begins with a convolutional layer followed by max pooling, then two additional convolutional layers, batch normalization, and ReLU with a skip connection that allows the input to bypass the signals from the previous layer if beneficial. This is followed by average pooling, and finally, a regression layer that computes a weighted sum of the averaged kernel outputs, where the weights are learned free parameters, along with a bias term to predict continuous gloss values.

In the first convolutional layer, the kernel size was 15 × 15 pixels for the one-layer model and 7 × 7 pixels for the three-layer model, determined through preliminary exploration of suitable architectures. The number of kernels varied between 1, 2, 4, and 9 for the one-layer model, and 9, 16, 32, and 64 for the three-layer model. One-layer networks were trained for 30 epochs, and three-layer networks for 90 epochs, with both trained until their performance reached a plateau. Training began with an initial learning rate of 0.01, which was reduced by a factor of 10 every 30 epochs. We used the Adam optimizer and mean absolute error as the evaluation function. Model training was conducted on a DGX A100 system (four NVIDIA A100 GPUs each with 80GB video memory, Dual AMD Rome 7742 CPU, 128 cores total, 2.25 GHz base) running Ubuntu 22.04 LTS. While each individual model was lightweight, the system enabled efficient parallel training of the many networks required for cross-validation.

### Training and validation procedure

Our evaluation procedure for the one-layer and three-layer models consists of two independent sets of cross-validation. In the first set, networks were trained on 33 lighting environments (3,564 images) and tested on 3 novel lighting environments (324 images). This process was repeated 12 times until all lighting environments were used as test sets. In the second set, networks had been trained on 33 shapes (3,564 images) and tested on the 3 novel lighting environments (324 images). Again, this had been repeated 12 times until all shapes were used as test sets. Thus, each network architecture underwent 24 unique training and testing pairs, resulting in 24 trained models. The correlation coefficient reported in **Figure 3** is the average value across these 24 networks, each computed from 324 test images. Luminance statistics models and specular reflection models were evaluated in the same way for consistency, although no fitting was involved for the luminance statistics models.

ResNet18 models (with or without additional training images) and three-layer models with additional training images were not evaluated using 24-fold cross-validation due to the substantial computational time required for training each model. Instead, for these models, one of the 24 training–test pairs was randomly selected, and performance was assessed using a single evaluation (for models plotted with orange symbols in **Figure 3**, the training set included additional images).

For each training session, the 3,564 images were augmented to 320,760 images through horizontal flipping and random size cropping, where a randomly selected sub-region (80–100% of the image) was rescaled to the network’s input size. This conservative augmentation approach was chosen because certain manipulations, such as vertical flipping, may disrupt the geometrical regularity in natural environments, potentially affecting the representativeness of gloss judgments.

## Acknowledgments

The authors thank Ana Serrano for providing the associated data for the rendered image dataset, and Matteo Toscani for providing the real-world photographs used to validate our models. We also thank Annika Zentel for her assistance with data collection in the laboratory-based experiment. TM was supported by a Sir Henry Wellcome Postdoctoral Fellowship from Wellcome Trust (218657/Z/19/Z). This research was also supported by “The Adaptive Mind,” funded by the Excellence Program of the Hessian Ministry of Higher Education, Science, Research, by the DFG SFB-TRR-135 “Cardinal Mechanisms of Perception” (project No. 222641018, TPs C1 and C2), and by a Marsden Fast Start Grant to KS from the Royal Society of New Zealand (project MFP-UOA2109), and European Research Council Advanced Grant Color3.0 (project number 884116) to KRG and European Research Council Advanced Grant STUFF (project number 101098225) to RWF. For the purpose of open access, the author has applied a CC BY public copyright license to any Author Accepted Manuscript version arising from this submission.

## Data availability

All behavioural data, stimulus images, model data are available on the GitHub page at https://github.com/takuma929/gloss_tinynetworks under non-restrictive MIT license.

Behavioural data and model data from the Serrano dataset, which were used to validate our models, are available at the following links: https://mig.mpi-inf.mpg.de/ (behavioural data) and https://github.com/Hans1984/material-illumination-geometry (model data).

## Code availability

All custom analysis codes used to reproduce figures in the manuscript are available on the GitHub page at https://github.com/takuma929/gloss_tinynetworks.

## Supplementary Information

### Gloss and the Ward reflectance model

In this study, we use the term *gloss* to refer to the subjective impression of shine or lustre that arises from the way surfaces reflect light. A complete physical description of a surface’s reflectance properties is described by the bidirectional reflectance distribution function (BRDF; [1]), which captures the amount of light reflected by the surface in every direction as a function of light arriving from every direction. Numerous low-parameter analytical approximations of typical surface BRDFs have been developed (see [2] for a review). Here, we employed the widely-used Ward BRDF model [3, 4] to generate our stimuli. In this model, the reflected light from a given point on an object’s surface is represented as a linear weighted sum of diffuse reflection, which primarily determines the body colour, and specular reflection, which gives rise to the sensation of gloss. **Figure S1** illustrates diffuse reflection (panel a) and specular reflection (panel b). Panel c shows an example object rendered with only diffuse reflection. In contrast, glossy objects additionally exhibit specular reflection that conveys direct information about the illuminant spectrum and thus about the light incident on the surface (panel d). In the experiment, we systematically varied the magnitude of the specular reflectance (i.e., the strength of the specular lobe). As shown in the main text, it is important to emphasize that the physical strength of the specular reflection does not directly correspond to perceived glossiness, but is correlated with it, much as perceived surface lightness is correlated with the surface’s diffuse albedo.

**Figure S1.**
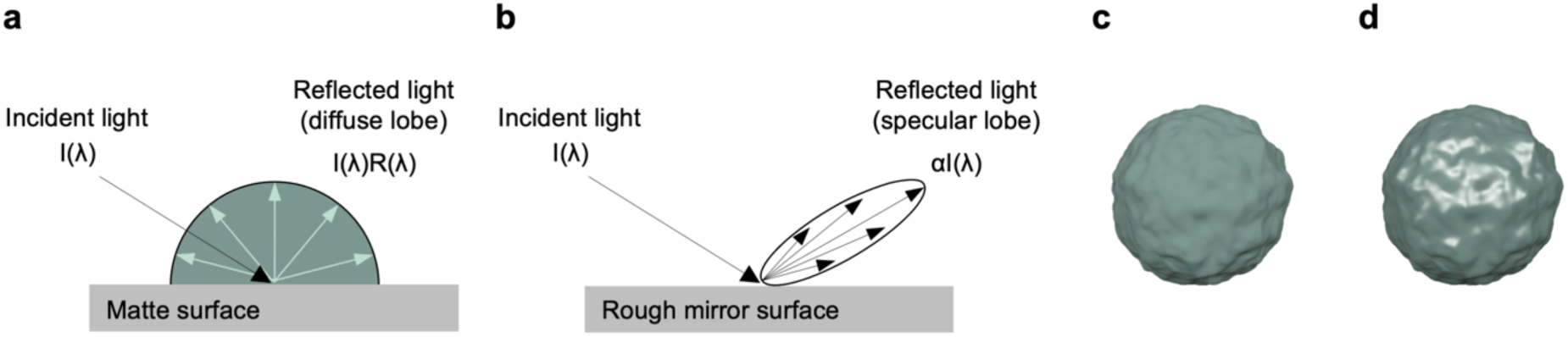
Two types of reflection—diffuse and specular—that occur when light interacts with an object. (a) Diffuse reflection, in which the reflected light’s spectral content is the product of the illuminant’s spectrum I( λ ) and the surface’s spectral reflectance R( λ ). (b) Specular reflection, which is a direct reflection of the incident light and therefore preserves the illuminant’s spectral content. (b) An example of a matte object exhibiting only diffuse reflection. (d) An example of a glossy object exhibiting both diffuse and specular reflections.

### Object geometries and lighting environments

To generate test images, we selected 36 three-dimensional meshes of everyday objects from Evermotion (https://evermotion.org) and 36 image-based illumination maps, as shown in **Figure S2 and S3**. For a reference image, we chose an Uffizi light probe, as in a previous study [5], and generated the geometry of a bumpy sphere using the 3D modeling software Blender (version 2.79b). These illumination maps and shapes were selected to cover a wide range of characteristics, with different complexities, which we thought would be key to identifying unique human behaviors that decorrelate from physical reflectances. Panel C shows the chromatic distribution of each lighting environment. There is a tendency for pixel colors to spread along the vertical blue-yellow axis, aligned with the daylight locus [6].

**Figure S2.**
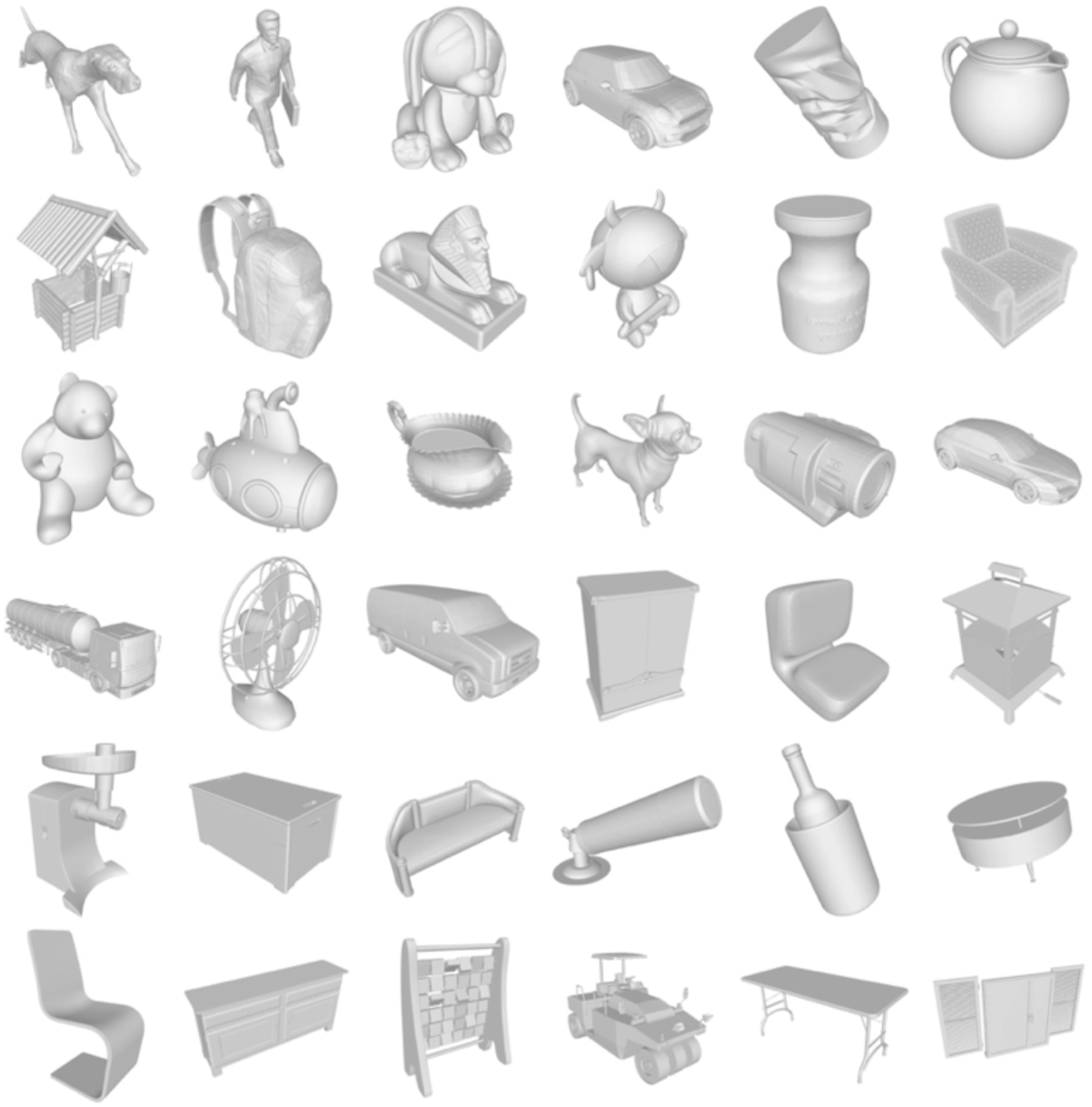
Thirty-six object geometries used to generate test images. They were selected to be as diverse as possible.

**Figure S3.**
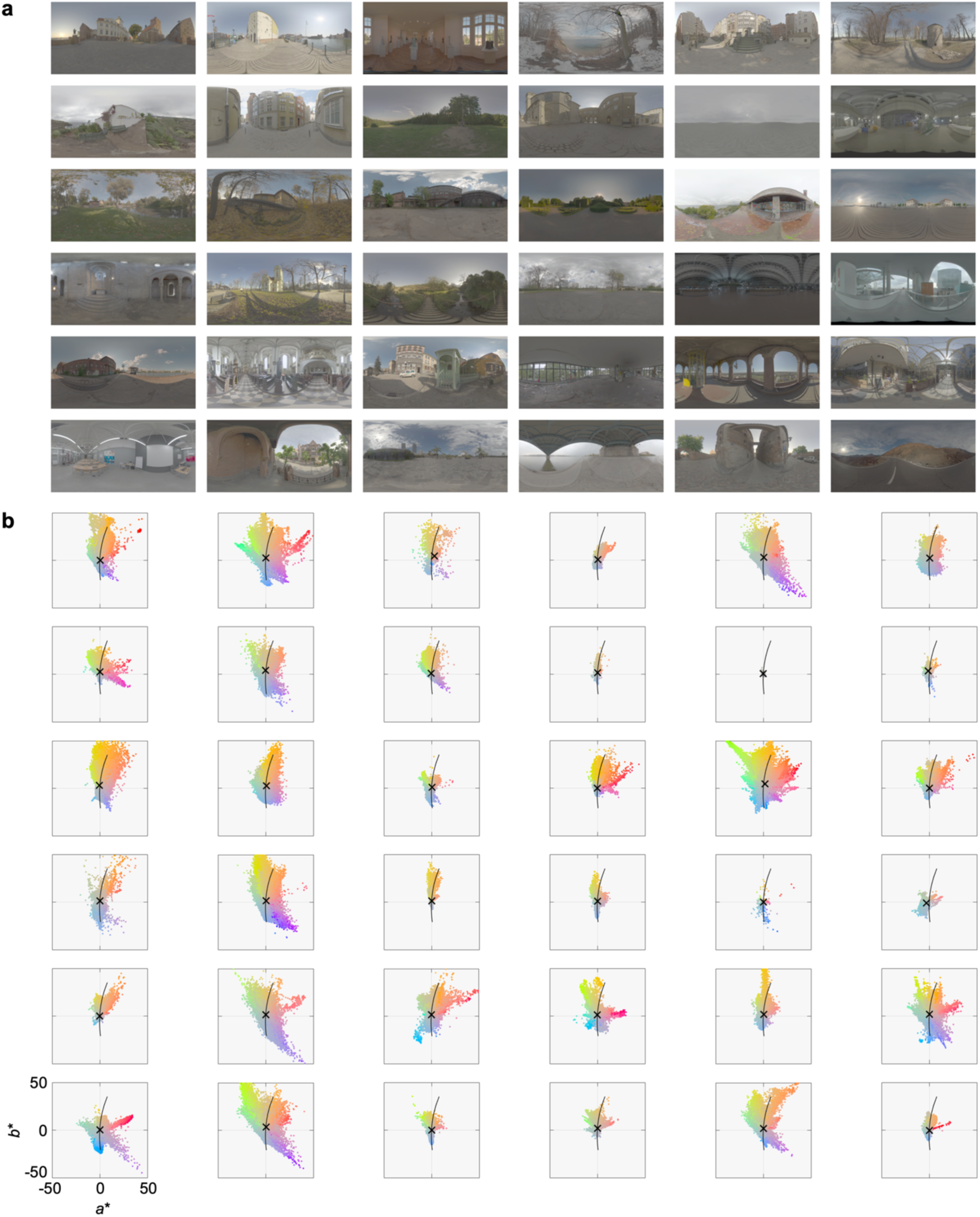
Lighting environments and their chromatic statistics. **a** Thirty six lighting environments. **b** Chromatic distribution of each lighting environment in L*a*b* color space. We randomly sampled 10% of pixels to reflect the density of distribution. The black cross symbols show mean chromaticity and the black curve shows the daylight locus. The white point was set to D65. The layout is the same as panel a.

### Observer settings for each lighting environment and shape

**Figures S4** and **S5** show mean observer settings compared to the physical ground-truth, grouped by each lighting environment and each shape, respectively. Each data point represents one image, with each plot containing 108 images (36 lighting environments or 36 shapes × 3 viewpoints). The aim of this analysis is to determine whether a specific lighting environment or shape consistently induces high or low gloss levels. For lighting environments, although correlations between human settings and physical ground-truth vary between 0.40 and 0.68, observer setting patterns were generally similar, with no noteworthy trends. In contrast, there are clearer differences across shapes. Shapes ranked high tend to be round, presumably because they generate highly visible specular reflections, while flat objects ranked low tend to produce low perceived gloss. This is visually intuitive, as specular highlights for the flat objects are widely spread on their surfaces and not easily visible. Overall, these findings show that when focusing on a particular shape or lighting environment, perceived gloss somewhat correlates with the underlying physical specular reflectance. It is the use of diverse shapes and lighting environments that decorrelates human responses from the physical ground-truth labels in our behavioural data.

**Figure S4.**
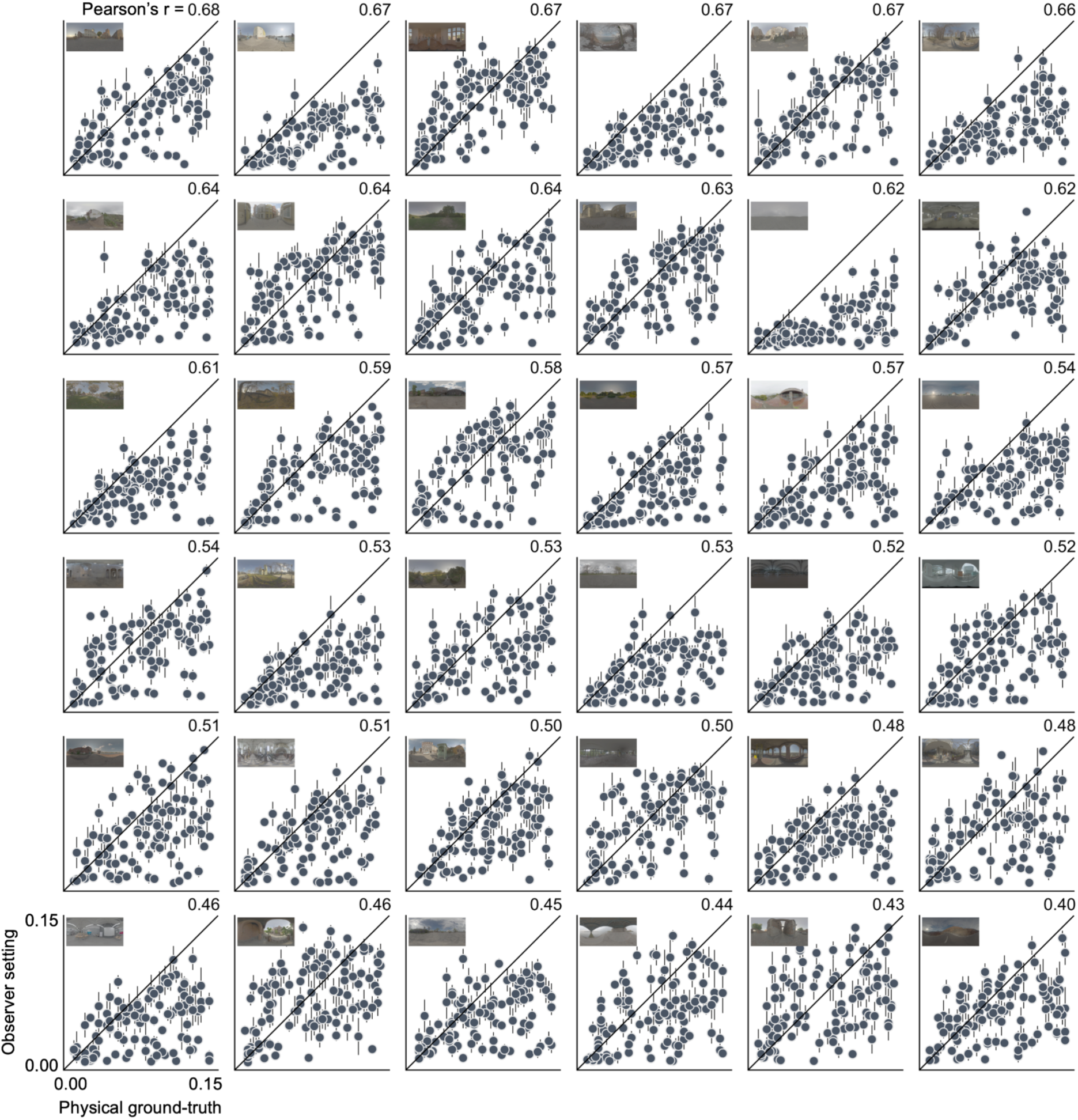
Average observer settings compared to physical ground-truth for images grouped by lighting environment. All images for each specific lighting environment were included in each panel, resulting in 108 data points (36 shapes × 3 viewpoints). The upper right value shows Pearson’s correlation coefficient, used to rank each panel. The vertical error bars represent ±S.E. across observers.

**Figure S5.**
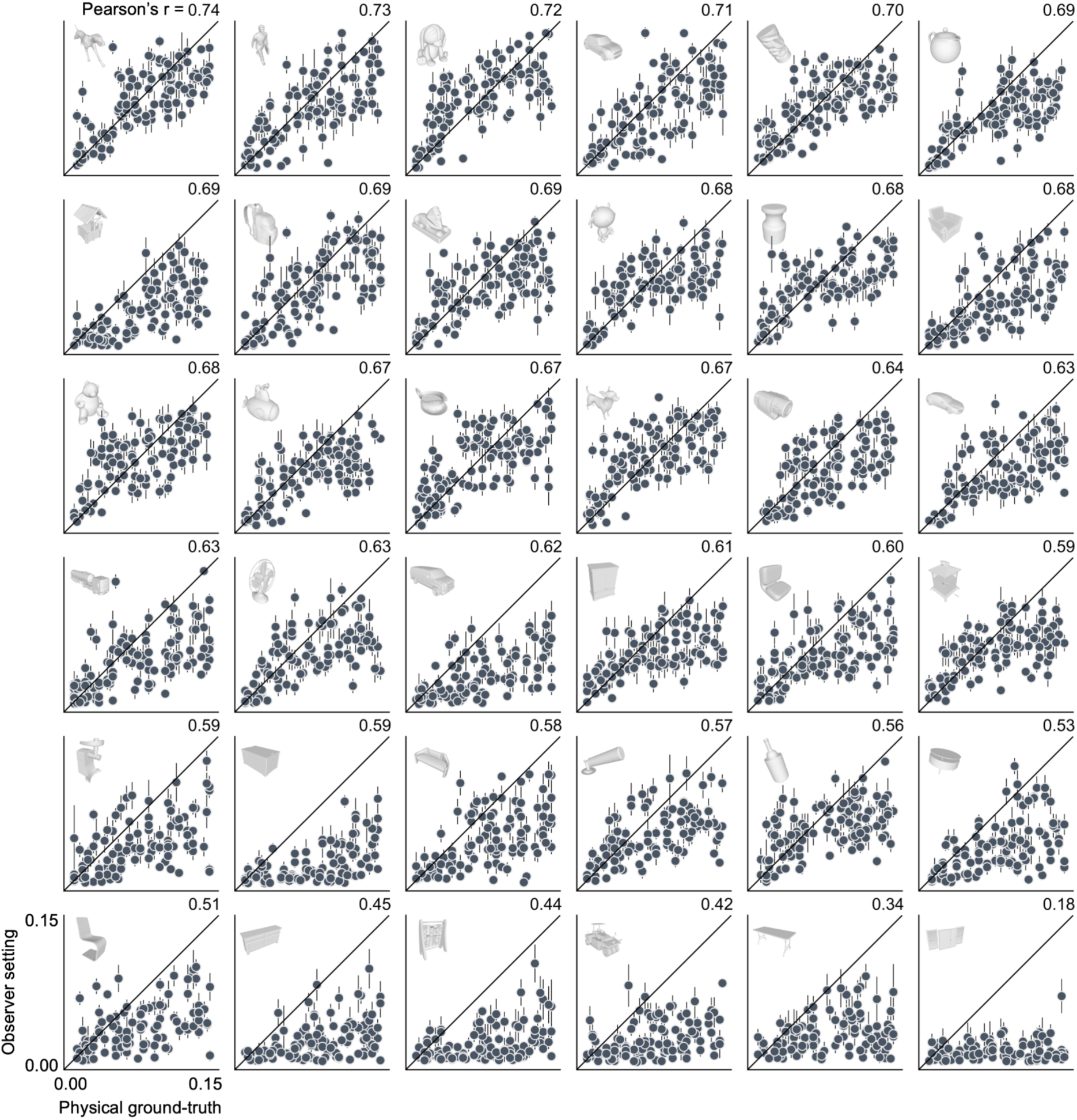
Average observer settings for images grouped by object shapes. Each plot contains 108 images (36 lighting environments × 3 viewpoints). The upper-right value indicates the Pearson correlation coefficient used to rank panels. Vertical error bars denote ±S.E. across observers.

### Laboratory experiment

#### Motivation

Our main experiment was conducted online to efficiently collect a large number of perceptual labels. However, there are general concerns with online experiments [7], such as the lack of monitor calibration, absence of an experimenter, and lack of in-person communication, which may affect task comprehension. To assess the quality of the online data, we conducted a validation experiment in a controlled laboratory environment as follows.

#### Observers and ethics

Twenty observers (13 females and 7 males) were recruited. Observers’ age ranged from 18 to 36 and mean ± standard deviation was 23.4±4.06 years. All observers had normal or corrected-to-normal visual acuity and normal color vision, screened using Ishihara Pseudo-isochromatic color plates [8]. Informed consent was obtained before the experiment. Before the experiment, the experimenter explained the task of the experiment to the observers. All procedures were approved by the local ethics committee at Justus Liebig University Giessen.

#### Test images

We used two of the image sets that had been shown in the main experiment to 21 and 23 online observers. Each set contained 72 images, excluding 12 common images. All lab-based observers judged all images from both sets.

#### Apparatus

Images were displayed on a 24-inch LCD monitor (ColorEdge CG2420, 1920×1200 pixels, frame rate 60 Hz; EIZO, Ishikawa, Japan) that allows for 10 bits per color channel (red, green, and blue). Gamma correction and spectral calibration were performed using measurements from a spectroradiometer (CS-2000; KONICA MINOLTA, Inc., Tokyo, Japan). The code to run the experiment was prepared in MATLAB using custom functions as well as functions provided in PsychToolbox-3 [9]. A chin rest was used to maintain the viewing distance.

#### Procedures

The experiment was conducted in a dark room. Observers placed their forehead and chin on the chinrest, to keep the viewing distance constant at 49 cm from the LCD monitor. Observers viewed the monitor binocularly. In each session observers completed 144 trials (72×2 images), and all observers completed 2 sessions, totaling 288 trials. The images were presented in a random order. Otherwise, the experimental procedure and the task of the observer were identical to the online experiment. Median response time was 6.18 sec per trial across — longer than in the online experiment (4.89 sec per trial).

#### Results

Panel **a** in **Figure S6** compares observer settings between laboratory and online experiments, where each data point represents the average across all observers. In both image sets, we found a near-perfect correlation (*r*(72) = 0.98, *p* = 2.1×10^-51^ for imageset 1; *r*(72) = 0.98, *p* = 9.3×10^-67^ for imageset 2). These very high correlations likely result from averaging data from a large number of observers. We also created a histogram of correlations for all possible pairs of individual observers (panel **b**). In this plot, we observed that the median correlation was higher than 0.8, which is not dramatically lower than the within-observer correlation shown in panel **c**. In summary, these comparisons demonstrate excellent consistency between online and laboratory-based experiments, suggesting that online data collection may be feasible for certain research questions and visual tasks.

**Figure S6.**
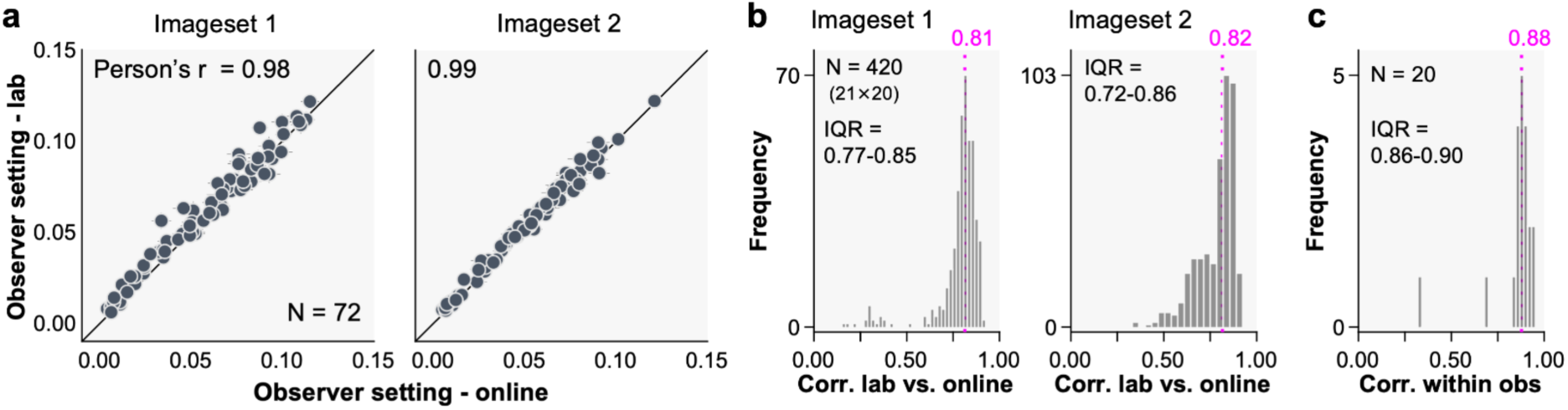
Comparison between online and offline results. **a** Laboratory experiment data plotted against online experiment data, with each data point representing the average across observers. **b** Histogram of correlations across individual observers between laboratory and online experiments. The median is shown in magenta. **c** Histogram of within-observer correlations computed between sessions 1 and 2 over 144 images.

### Extended analysis of blob-and-ridge kernels

This section extends the analysis presented in **Figure 4** of the main text. To compare kernels developed in human-like single-kernel models with those that emerged in the physical ground-truth networks, **Figure S7a** shows the single kernels obtained from 24 physical ground-truth networks. Diagonal ridges are absent in these kernels, unlike in the human-like networks, highlighting their importance as a feature specific to human gloss judgments. Furthermore, the kernels from the physical ground-truth networks exhibit higher luminance contrast than those from the human-like networks. This suggests that capturing the physical ground truth requires encoding features spanning a broader luminance range, whereas the kernels to predict human perception appear to compress this dynamic range. This finding is consistent with Pellacini’s c parameter being the cube root of specularity in the Ward reflectance model [10].

To further examine the role of projected object geometry in the training images, we rotated the objects around the X rotation axis by 45° and 90° and generated 3,888 object images with otherwise identical rendering parameters. We then trained a single-kernel network under the assumption that human gloss judgments would remain unchanged. As a result, the orientation tuning of the emergent kernel also shifted (**panel b**) to 17° and 161°.

As shown in **panel c**, we next rotated the lighting environments in two ways: first, by rotating the light probe on the image plane by 45° and 90° so that horizontal structures became diagonal and vertical; and second, by changing the elevation by 45° and 90°. The in-plane rotations had only limited effects— the 90° rotation did not alter kernel orientation, and the 45° rotation produced only a slight ridge shift to 61°. Similarly, the elevation rotations had no noteworthy impact.

In sum, this analysis suggests that the orientation of the learned kernel ridge is influenced more strongly by the projected surface geometry in the image set than by the directionality of the lighting in the light probe. In our stimulus generation process, objects generally maintained an upright posture (with slight variations in tilt of ±15° rotation along X and Y rotation axes). Although the object shapes and poses varied in our dataset, it is possible that many 3D models—whether artist-designed or everyday objects—share geometric regularities that naturally produce oriented highlights around 45° and 135°, thereby influencing the orientation selectivity of the learned kernels. The extent to which such regularities reflect shape variations in the real world remains an interesting empirical question beyond the scope of the present study.

**Figure S7.**
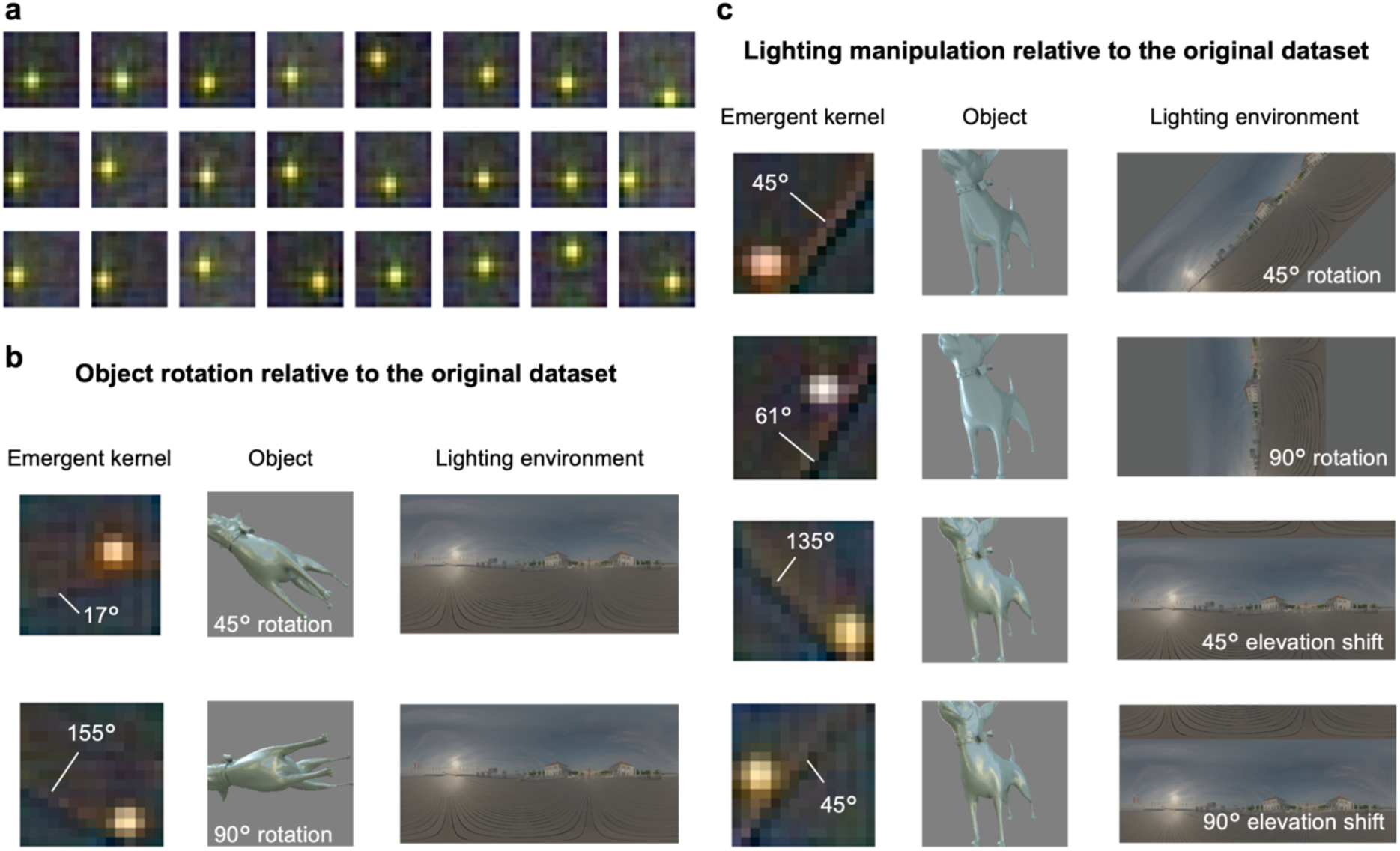
Extended analysis of “blob-and-ridge” kernel. **a** Twenty-four emergent kernels from single-kernel models trained on physical ground-truth labels. **b** Effects of object rotation (45° and 90°) on the emergent kernel. **c** Effects of lighting manipulations (in-plane rotation and elevation shift) on the emergent kernel.

This suggests that the presence of the oriented ridges in the kernels, really is something to do with human perception, rather than just being a generically useful feature for estimating reflectance in our training images. This likely reflects the fact that the human visual system is exposed to a much more diverse visual diet, and has to solve many more visual tasks than just estimating specular reflectance. Presumably, when asked to identify gloss, humans rely especially on features that distinguish specularity from other sources of image contrast (see previous point 3).

### CNN-based models that take the surrounding context information into account

We removed the surrounding context from each image, set background pixels to mid-gray (R=G=B=128), and used the resulting images to train the CNN-based models presented in the main text. This decision was made because our shallow network model lacks the computational capacity to effectively segregate the background from the object while also predicting gloss levels. However, the surrounding context might contain useful information that could help CNNs better predict human responses. To explore this possibility, we added an additional stream to our existing models, incorporating information from the surrounding region into the regression layer at the end to determine gloss levels.

**Figure S8** illustrates the architectures of such models and their performance in predicting human responses for the one-layer model (panel **a**) and the three-layer model (panel **b**). The results indicate that including context information in the network stream results in no performance improvement for both models (two-tailed paired t-test; *t*(23)=0.57, *p* = 0.57, Cohen’s *d* = −0.12 for one-layer model; *t*(23)=0.55, *p* = 0.588, Cohen’s *d* = 0.11 for three-layer model), suggesting that the surrounding context does not provide significant additional information. It is possible that useful cues are present but not captured by the networks. Nevertheless, this observation aligns with past empirical evidence that observers do not rely on surrounding contextual information when judging surface glossiness [11].

**Figure S8.**
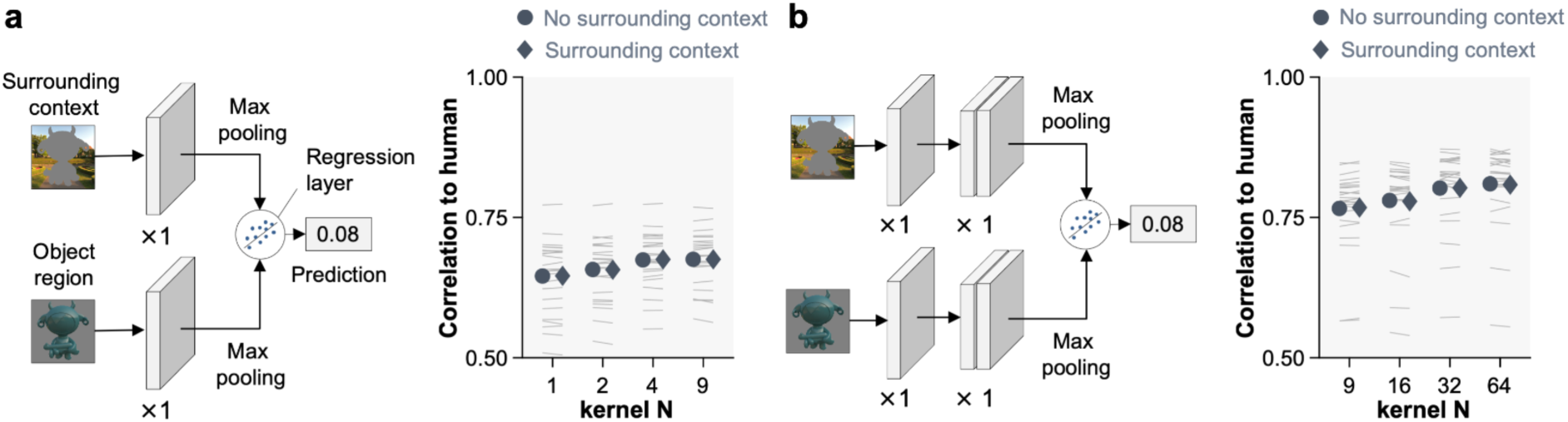
Testing whether surrounding context provides useful information for predicting human responses. **a** The left network illustrates the architecture of a one-layer model with an additional stream for processing surrounding context. The circles indicate the correlation coefficient with human responses, averaged across 24 networks when no surrounding context was used, as reported in the main text (Figure 3). The diamonds show the results when surrounding context was included. **b** The same analysis for a three-layer network. Lines indicate performance changes for each of 24 networks. In both cases, including surrounding context information resulted in little performance improvement.

#### Textured objects

In our main image set, all object surfaces had uniform base colors. However, real-world surfaces can sometimes have high-contrast textures, which our filter-based model may misinterpret as specular reflections.

Thus, as shown in **Figure S9a**, we tested the single-kernel model’s response to objects from a subset of the original dataset (1,296 images) with added surface textures. When the texture had the same luminance as the surrounding body color and introduced only chromatic contrast, the predicted gloss level remained stable (panel **b**, left). However, when the texture’s luminance was increased to 1.5 times that of the body color, the model often misinterpreted it as a specular highlight, leading to a systematic increase in predicted glossiness (panel **b**, right).

However, if textured objects had been included in our training data, the model could have learned to suppress responses to texture and become more selectively tuned to specular highlights. To test this, we retrained the model using both textured and non-textured objects. Half of the training images were textured and the other half were non-textured, but pairs of the same object (textured and non-textured versions) were not included together. Since the textured versions lacked their own labels, we assigned them the same gloss labels as their non-textured counterparts of the same object. We then examined the filters that emerged (panel c, upper). In the two-kernel model, one excitatory kernel similar to the original emerged, along with a second inhibitory kernel (panel **c**, lower). The model predicts glossiness as a weighted sum of the responses from these two filters. Notably, the inhibitory kernel showed strong selectivity to texture (Figure 9d), effectively functioning as a texture-canceling filter that counteracts misclassification driven by texture contrast. We found that the two-filter network suppressed the systematic increase in predicted glossiness for textured surfaces (panel e), with the mean absolute error (MAE) significantly lower than that of the single-kernel model (one-sided Wilcoxon signed-rank test, *z*(1295) = –9.13, *p* = 3.52×10⁻², effect size *r* = –0.25). Normality of the difference scores was not confirmed (Lilliefors test, *p* < 0.001). Our high-contrast textures included sharp spatial variations in both chromaticity and luminance. The colorful suppression kernel that emerged is specifically tuned to respond strongly to spatial chromatic variation, such as that found in textures. In contrast, specular highlights typically appear as smooth luminance gradients in low-saturation yellow-blue or white regions, which elicit only a weak response from this kernel. The network improves its selectivity for specular highlights by offsetting texture-driven activation in the excitatory kernel with the inhibitory kernel’s response.

Thus, while texture poses a challenge, our results suggest that this issue can be mitigated by incorporating additional filters and exposure to textured objects in the training set. Naturally, if the texture exhibits spatial structures that more closely resemble specular highlights, suppression becomes more difficult—not only for our model, but likely for any model, and potentially even for human observers. Nonetheless, this illustrates how a data-driven approach can uncover a concrete computational solution to a long-standing challenge in the field. In real-world viewing, motion and binocular cues may further assist in distinguishing surface texture from specular highlights by revealing differences in their relative motion or disparity patterns [12, 13].

**Figure S9.**
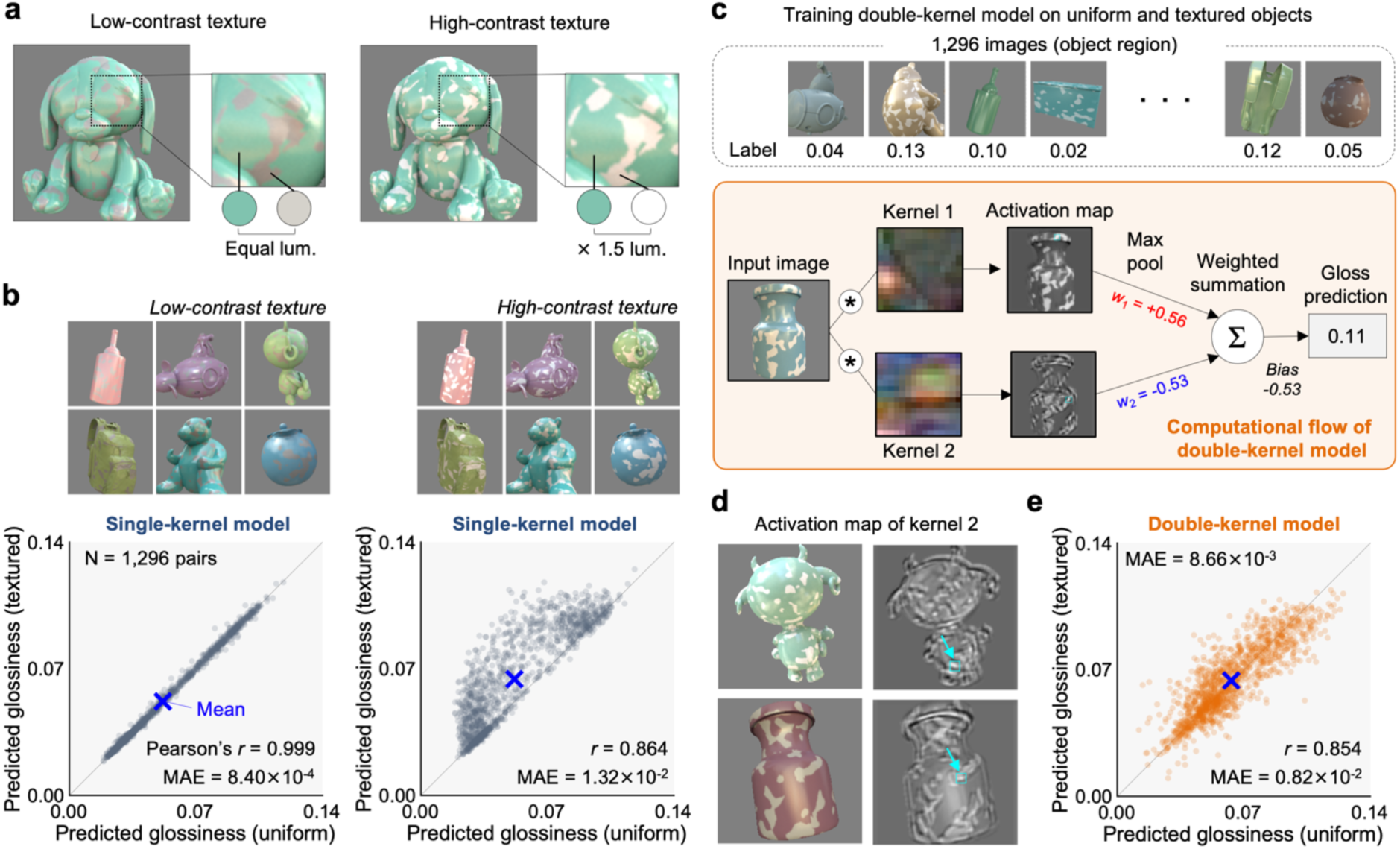
Evaluation of model responses to textured objects. **a** Two types of textures were applied to a subset of the original dataset: low- and high-contrast textures, each with either equal luminance or 1.5 times higher luminance relative to the body color. **b** Model responses to the original image set (x-axis) versus the textured image set (y-axis), shown separately for low-contrast textures (left) and high-contrast textures (right). **c** A model with two kernels was trained on a new dataset containing both uniform and textured objects. The model convolves the input image with two kernels, applies max pooling to the activation maps, and predicts glossiness by taking a weighted sum of the max-pooled values plus a bias term. **d** Example activation maps of the inhibitory kernel. The cyan square highlighted by an arrow indicates the image region that produced the highest activation in the double-kernel model. **e** The double-kernel model reduces the increase in predicted glossiness caused by texture patterning.

